# Decoding Cellular Stress States for Toxicology Using Single-Cell Transcriptomics

**DOI:** 10.1101/2025.06.10.657506

**Authors:** Imran Shah, David Gallegos, Brian Robinette, Bryant A. Chambers, Dennis J. Eastburn, Douglas A. Bell, Michelle R. Campbell, Suzanne N. Martos, Salvatore Camiolo, Kevin S. White, Nicole Martin, Gioele Montis, Joel McComb, Bruce Seligmann, Brian N. Chorley

**Author notes:** Special Volunteer (retired), National Institute of Environmental Health Sciences - NIH. **Correspondence to be addressed to:** Imran Shah Center for Computational Toxicology and Exposure (CCTE) Office of Research and Development U.S. Environmental Protection Agency 109 TW Alexander Drive (D130A) Research Triangle Park, NC 27711.

## Abstract

We applied the TempO-LINC® platform to generate single-cell transcriptomic (SCTr) profiles of ∼40,000 HepaRG cells exposed to etoposide, brefeldin A, cycloheximide, rotenone, tBHQ, troglitazone, and tunicamycin at three concentrations for 24 hours. SCTr enabled a detailed analysis of adaptive stress response pathways (SRPs), including the unfolded protein response (UPR), oxidative stress response (OSR), heat shock response (HSR), and DNA damage response (DDR). Troglitazone upregulated lipid metabolism genes (*PLIN2*, *ACOX1*) along with HSR and UPR activation, with co-expression of *DNAJA1*, *HSP90AA1*, and *DDIT3* in subsets of cells. Brefeldin A and tunicamycin strongly induced UPR markers (*HSPA5*, *SYVN1*, *LMF2*, *PDIA4*) in subsets of cells, with some also expressing apoptotic (*DDIT3*, *CASP8*) and autophagic (*SQSTM1*) genes, indicating diverse stress responses. Rotenone activated *GDF15*, *TRIB3*, and *DDIT3* in a fraction of cells, accompanied by *PLIN2* and mild UPR induction, reflecting heterogeneous mitochondrial stress responses. We scored individual cells using literature-derived SRP gene signatures to characterize overall stress phenotypes and clustered them using a generalized Jaccard metric. The clustering revealed five phenotypic groups spanning cell states associated with homeostasis, adaptive responses, terminal outcomes, autophagy, and apoptosis. By systematically analyzing the distributions of cells in different states across treatments, we visualized dynamic shifts in cellular subpopulations responding to chemicals, revealing early stress responses and potential transitions to cell death. Our findings suggest the utility of SCTr in decoding stress states that could provide possible insights into transitions between cellular adaptive and terminal transitions involved in toxicity.

## Introduction

Traditional animal-based toxicity testing remains too resource-intensive and time-consuming to support the comprehensive safety evaluation of the vast number of chemicals currently in commerce (US EPA, 2019). In response, the U.S. Environmental Protection Agency (USEPA) and international stakeholders have prioritized the development of New Approach Methodologies (NAMs) that reduce animal use while improving human relevance and mechanistic insight (USEPA, 2018; Thomas et al., 2019). High-throughput transcriptomics (HTTr) is a key NAM that enables rapid, genome-wide profiling of chemical-induced gene expression changes in human cell models, supporting both hazard identification and mechanistic interpretation (Bundy et al., 2024; Harrill et al., 2024). However, conventional HTTr approaches typically rely on bulk RNA sequencing, averaging gene expression across all cells in a population and obscuring heterogeneity in cellular responses (Potter 2018) that may be critical to understanding toxicity (J. Chen et al. 2024).

Single-cell transcriptomics (SCTr) offers a powerful approach for resolving heterogeneous cell states that may underlie differential sensitivity to toxicants and play key roles in pathology and disease (Aldridge and Teichmann, 2020). Unlike bulk transcriptomics, which captures average responses across cell populations, SCTr can uncover rare or divergent cellular phenotypes critical for understanding mechanism-based toxicity. This resolution makes SCTr a promising complement to HTTr, with the potential to identify unique stress-induced biomarkers and mechanistic signals that may be masked in population-level data.

We are particularly interested in using SCTr to explore cellular tipping points, previously identified across multiple cell types and assay platforms (Shah et al. 2016; Frank et al. 2018; Saili et al. 2020). These tipping points represent critical thresholds beyond which stress responses shift from homeostatic adaptation to irreversible terminal cellular outcomes, such as apoptosis or autophagy (Fulda et al., 2010). By characterizing adaptive and terminal cell states, SCTr could elucidate the molecular switches that underlie transitions to toxicity and allow for the identification of early warning markers before overt cellular damage occurs (Szegezdi et al. 2006). Such markers could improve confidence in HTTr-based NAMs by clarifying when chemical-induced gene expression changes are likely to reflect protective responses versus early indicators of pathology. This mechanistic resolution is critical for improving the predictive power of and confidence in in vitro models in chemical risk assessment (Jennings 2015).

Adaptive stress response pathways (SRPs) are evolutionarily conserved cellular programs that maintain homeostasis under conditions of chemical or environmental stress. These include the unfolded protein response (UPR), oxidative stress response (OSR), heat shock response (HSR), and DNA damage response (DDR), among others (Hiemstra et al., 2017; Vihervaara and Sistonen, 2014). While SRPs are protective under transient stress, prolonged activation can result in terminal outcomes such as apoptosis, autophagy, or senescence—cellular fates that have been implicated in chemical-induced liver diseases, including steatohepatitis and hepatocellular carcinoma (Kanda et al. 2018; Spaan et al. 2019; Kouroumalis et al. 2021). Thus, distinguishing adaptive versus adverse responses is a critical need in implementing NAMs for regulatory toxicology (Middleton et al., 2017). Mapping cell states along this continuum may help relate molecular changes to phenotypic outcomes and improve the interpretation of *in vitro* assays in terms of potential *in vivo* pathology.

In this preliminary study, we evaluated whether SCTr could resolve stress-related cellular phenotypes and their progression under chemical stress. We used the TempO-LINC® platform to profile ∼40,000 single HepaRG cells exposed to seven well-characterized reference stressors—etoposide, brefeldin A, cycloheximide, rotenone, tBHQ, troglitazone, and tunicamycin—each applied at three concentrations for 24 hours. HepaRG cells were selected due to their ability to differentiate into hepatocyte- and cholangiocyte-like cells and their extensive use in liver toxicity research (Guillouzo et al., 2007; Gerets et al., 2012). These exposures were chosen to activate SRPs relevant to liver toxicology, to identify chemically induced stress states.

To this end, we analyzed stress pathway activities and downstream cell fate markers, including apoptosis, autophagy, and senescence. Literature-derived gene signatures and pathway activity scores were used to cluster individual cells into phenotypic groups, enabling a systems-level view of how cells transition from homeostasis through adaptation and, in some cases, to terminal outcomes. By capturing this heterogeneity and putative responses, we demonstrate the potential of SCTr to provide mechanistic insights into early toxicity events and cellular tipping points—key features that can enhance the predictive value of NAMs and support their application in chemical safety assessment.

## Methods

The workflow for the entire study is outlined in Figure 1, and the steps are described below.

**Figure 1.**
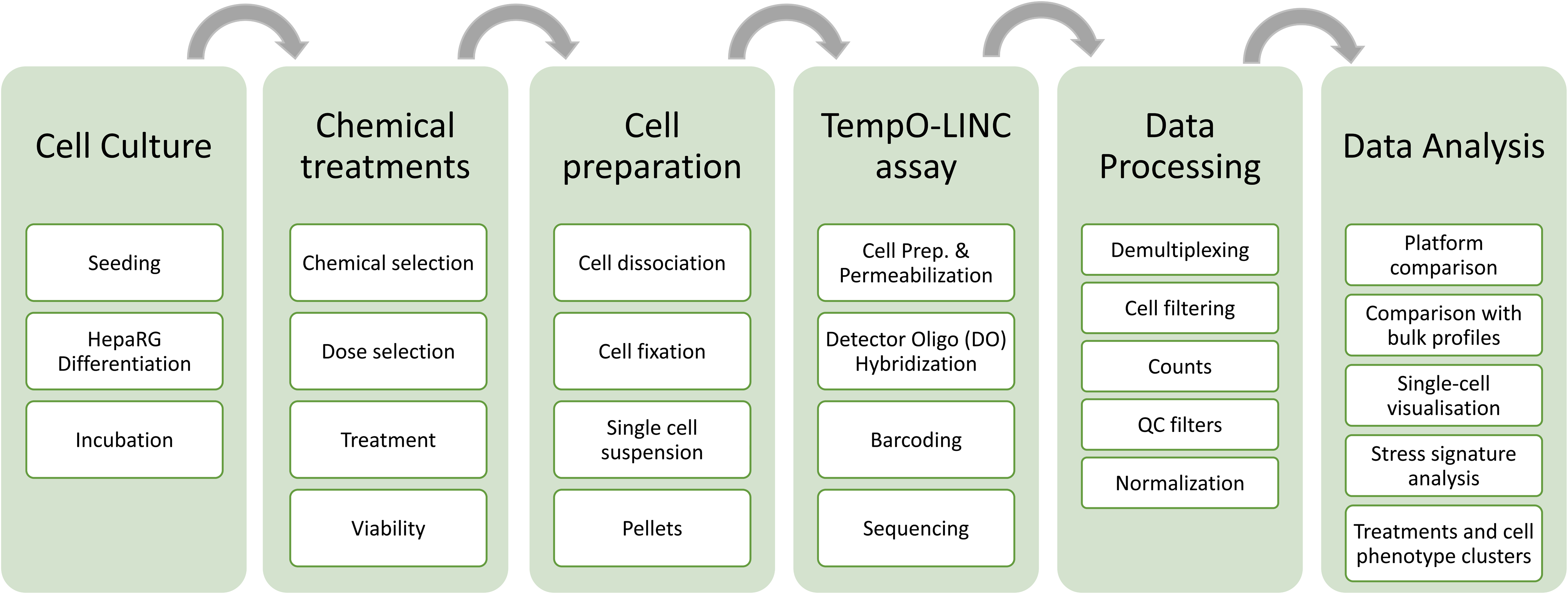
Single-Cell transcriptomics workflow. This figure illustrates this study’s overall workflow for conducting single-cell transcriptomics (SCTr). The process begins with cell culture, followed by chemical treatments to induce cell stress responses. Cells are then prepared for single-cell analysis by fixation, dissociation, suspension, and pellet formation. The TempO-LINC assays are then used to generate the SCTr data. Finally, the data analysis stage involves identifying gene expression changes, clustering cells, and performing connectivity mapping to assess pathway activation in response to treatments, and identifying phenotypic clusters of cells.

### Chemical selection

A set of 8 chemicals (including DMSO control) was tested to investigate distinct mechanisms of cellular stress responses (Table 1). Chemical concentrations were selected to ensure that they did not induce significant cell death in HepaRG cells by keeping them below the AC50 for cytotoxicity, which was assessed by measuring LDH release. Brefeldin A, recognized for its role in inducing the UPR by disrupting ER-to-Golgi protein transport (Sciaky et al. 1997), was used at concentrations of 0.001, 0.01, and 0.100 µM. Etoposide, a well-known DDR activator and a topoisomerase II inhibitor (Hande 1998), was included at doses of 0.25, 2.5, and 25 µM to assess DDR activation and cytotoxicity. For investigating oxidative stress response (OSR), tert-butylhydroquinone was chosen for its ability to activate the *NRF2* pathway, a key player in cellular defense against oxidative stress (Kensler, Wakabayashi, and Biswal 2007), and was tested at concentrations of 3, 30, and 300 µM. Rotenone, known to induce mitochondrial stress by inhibiting the mitochondrial electron transport chain (Isenberg and Klaunig 2000), was tested at concentrations of 0.2, 0.4, and 0.8 µM. Cycloheximide, a protein synthesis inhibitor impacting many cellular processes (Schneider-Poetsch et al. 2010), was tested at concentrations of 0.200, 2.000, and 20.000 µM. Tunicamycin, another UPR inducer known for inhibiting N-linked glycosylation (Heifetz and Lennarz 1979), was tested at concentrations of 0.001, 0.010, and 0.100 µM. Lastly, troglitazone, a *PPARγ* agonist influencing lipid metabolism and mitochondrial function (Rogue et al. 2010) and can modulate PPARα pathway activity (Aljada et al. 2001), was tested at concentrations of 0.001, 0.010, and 0.100 µM. Lastly, 0.1% dimethyl sulfoxide (DMSO) was used as the vehicle control for all treatments. These test chemicals, spanning a range of concentrations, were selected to provide a comprehensive view of transcriptional responses across different cellular stress pathways without reaching a concentration level of each producing significant cell loss.

**Table 1.**
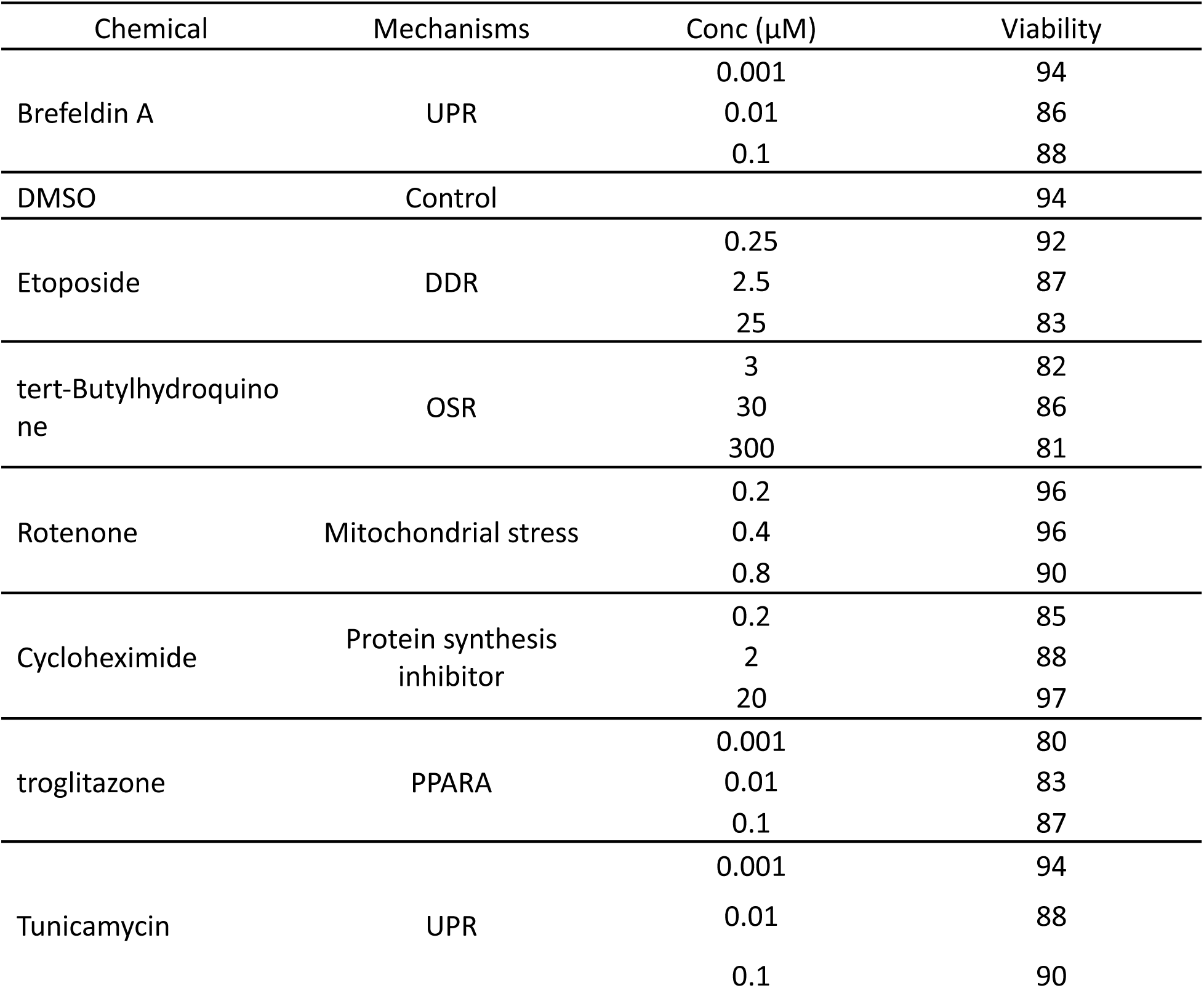
Chemical treatments, mechanisms, and viability.

### Cell culture and treatments

Terminally differentiated human hepatic cells (NoSpin HepaRG™) provided directly from the distributor (Lonza, Morrisville, NC) were seeded at 3X10^3^ cells/cm^2^ in 24-well plates coated with rat tail collagen (Sigma-Aldrich, Saint Louis, MO) at a density of approximately 5 µg/cm^2^. After dispersion and seeding, cells were allowed to settle in the plate at room temperature for ∼2 hrs in a biosafety cabinet before being transferred to an incubator set at 37°C, 5% CO_2_. Cells were incubated for 7 days in HepaRG Base Medium with Supplement (MH100; Lonza) plus Thawing and Plating additive (MHTAP; Lonza) before treatments, changing the media every 48-72 hrs (Angrish et al. 2017). Cell culture was observed for abnormal morphological changes or cell loss during this incubation by light microscopy. The observed morphology was clusters of cells described as islands of hepatocyte-like cells surrounded by cholangiocyte-like cells (Marion, Hantz, and Durantel 2010). During cellular treatment, 0.1% DMSO was used as a vehicle control, corresponding to the final vehicle percentage for all solubilized chemicals. Treatments were carried out for 24 h before termination. Given the high reproducibility of transcriptomic responses previously reported across independent HepaRG cultures, and in light of the exploratory nature of this study, we used a single treated culture per condition as a reasonable approach. This design enables the capture of representative cell population responses at single-cell resolution that could provide valuable insights into the heterogeneity and pathway-level activity induced by chemical exposure.

### Cell preparation: dissociation & fixation

Following treatment, cells were dissociated into a single-cell suspension. Cell counts and viability were determined with a fluorometric Nexcelom Cellometer automated cell counter (Nexcelom Biosciences, Lawrence, MA) using Acridine Orange (AO)/Propidium Iodide (PI) stain (Perkin Elmer, Inc., Waltham, MA), following manufacturer’s instructions, and then the cells were fixed and processed for single-cell library construction. Cells were dissociated in a 24-well plate for the TempO-LINC assay to provide the necessary cell input. Cells were initially incubated for 11 min at 37°C in 0.25% Trypsin-EDTA (Thermo Fisher Scientific) with 40 mM DTT, 0.1 mg/ml actinomycin D, 1 mg/ml anisomycin, and 6 mM RNaseOUT recombinant ribonuclease inhibitor (Thermo Fisher Scientific, Waltham, MA) in 250 µl volume, followed by an additional 15 min 37°C incubation after adding 250 µl 0.5% Collagenase Type 1 (Worthington, Lakewood, NJ) in EBSS (Earle’s Balanced Salt Solution) and DTT (Dithiothreitol) 40mM DTT (Dithiothreitol) and 6mM RNaseOUT. Using a wide-bore pipette tip, cells were then dissociated by gentle pipetting for 10 min, 2 ml of 37°C HepaRG Base Medium was added to stop enzymatic activity, and cells were pelleted at 200 xg for 3 min at 4°C. After media removal, cells were resuspended in 1 ml cell suspension media (1X PBS, 0.5% Opti-Prep Density Gradient Medium (Sigma-Aldrich), 0.1% Bovine Serum Albumin Fraction V (Thermo Fisher Scientific, Waltham, MA), 6 mM RNaseOUT) pre-chilled to 4°C. The cells were pelleted twice more at 200 xg for 3 min at 4°C, washing each time with fresh Cell Suspension Media. After the final wash, cells were filtered through a 40 µm nylon cell strainer, transferred, and stored in a Lo-Bind tube (Eppendorf North America, Hauppauge, NY).

For the TempO-LINC® assay samples, dissociated cells were fixed in 1% paraformaldehyde solution in 1 X PBS at room temp for 25 min with slow rocking, then quenched in 1:1 volume of 10 mM Tris 0.1% Pluronic F-68 (Thermo Fisher Scientific). Cells were then pelleted at 500xg for 5 min at 4°C and washed in 0.1% Pluronic F-68, 0.1% BSA Bovine Serum Albumin Fraction V, 0.1 U/µl Superase*In (Thermo Fisher Scientific) and 0.1 U/µl Enzymatics (Qiagen, Germantown, MD) ribonuclease inhibitors, and 40 mM DTT. After a final spin, most of the supernatant was removed, and cells were frozen at -20°C overnight, then stored at -80°C until library prep. For 10X chromium assay and Parse single-cell whole transcriptome analysis, cells were processed using the protocol and kit reagents provided with each platform (see Supplementary Information).

### Single Cell Transcriptomic Sequencing with TempO-LINC

TempO-LINC® single-cell sequencing libraries were prepared using the TempO-LINC v1.0 protocol (BioSpyder, Carlsbad, CA) by BioSpyder. Briefly, saponin-permeabilized cells from each sample were first hybridized with BioSpyder TempO-Seq whole transcriptome WT v2.1 Detector Oligos (DOs) that had been 5’ phosphorylated. Following hybridization, excess DOs were washed away, and the remaining DOs hybridized to target mRNA were ligated at 37°C for 1 hr. Samples were then labeled by distributing them into 48 wells of a microplate. Each well contained fixed cells from one of the treatment samples and one of 48 unique molecular barcodes (“barcode 1”). Then the barcodes were ligated to the 5’ phosphorylated detector oligos, still hybridized to mRNA within the fixed cells. This first barcode not only comprised a portion of each cell-specific barcode, but since each treatment sample was placed into a unique well for ligating on barcode 1, barcode 1 also identified the treatment condition. Cells were pooled following the first sample barcode ligation and then redistributed into wells of a plate containing 48 additional barcodes (“barcode 2”) were ligated onto barcode 1. Following pooling and a third round of barcode addition by ligation (“barcode 3”), after which unique molecular indexes (UMIs) were also added, cells were pooled once again and then distributed into 12 PCR tubes, and the fully barcoded DOs were amplified with indexed primers, a separate unique index being added to each PCR tube. Combining these barcodes and UMIs allows single-cell RNA sequencing to efficiently process large numbers of cells from multiple samples, allowing for comprehensive analysis of cellular heterogeneity and gene expression patterns (Eastburn et al. 2024). Amplified DNA was pooled, purified, and quantified before paired-end sequencing on an Illumina NovaSeq 6000 using an S4 flow cell. Sample library preps were verified and quantified using a Bioanalyzer by assessing fragment size distribution and ensuring the absence of primer dimers or degraded RNA.

### Raw data processing

To generate the TempO-LINC® single-cell expression data, barcoded reads were demultiplexed using software available from BioSpyder, with additional details provided elsewhere (Eastburn et al. 2024). Briefly, reads were scanned to find the sequences of the barcodes, allowing an offset of up to 3 nucleotides from their expected positions and 1 mismatch from their expected sequence to allow for gaps and variations due to minor sequencing errors. Reads featuring the same barcode combination were pooled together into barcode-associated clusters. The demultiplexed reads were then grouped by treatments into fastq files and made available at the National Center for Biotechnology Information (NCBI) Sequence Read Archive (SRA) under accession SRAXXXXX. A cell calling read threshold was set at 10,000 reads/cell based on inflection point analysis of a ranked barcode distribution plot (knee plot). Cell-associated reads were then aligned to the reference human whole transcriptome TempO-Seq probe sequences using bwa v. 0.7.17 (mem algorithm, parameters: -v 1 -c 2 -L 100). UMIs were used to collapse the reads originating from the same transcript and remove any PCR duplicates. The probes featuring a UMI count >0 for each single-cell alignment indicated an expressed/detected gene. Cell count tables were generated using the software featureCount v 2.0.1, including 33742 cells and 22,533 mRNA probes (BioSpyder human transcriptome probes v2.1). The count data for cells are made available in the NCBI Gene Expression Omnibus (GEO) under accession GSE299113.

### Data normalization

The raw cell count tables were filtered to include only cells with more than 10,000 counts and 1,000 detected genes. This ensured the inclusion of cells with sufficient transcriptomic information and depth, thereby enhancing the reliability and interpretability of the data (Lun, McCarthy, and Marioni 2016). The counts within each cell were normalized to 10,000 counts per cell to mitigate variations in gene expression levels due to differences in cell size or sequencing depth (Wolf, Angerer, and Theis 2018). Following normalization, the data were log_2_ transformed to stabilize the variance across the range of expression levels (Soneson and Robinson 2018). We applied batch correction to mitigate the impact of separate experimental runs using Harmony, which projects cells into a shared embedding by iteratively adjusting for known sources of variation, while preserving biological differences across cells <u>(Korsunsky et al. 2019)</u>. To focus on the most informative features of the dataset for visualisation, we identified the 2000 most highly variable genes employing the ‘seurat_v3’ algorithm, which selected genes with significant variability in expression. These genes often drive biological processes and cellular heterogeneity (Stuart et al. 2019). To facilitate visualization, we performed stratified sampling to create a balanced subset of cells with equal representation from each treatment condition. This balanced dataset was then visualized in two dimensions using t-distributed stochastic neighbor embedding (t-SNE) with perplexity = 25 and learning rate = 1000.

### Platform comparison

A platform comparison was conducted to benchmark the performance of three single-cell transcriptomics (SCTr) technologies: BioSpyder TempO-LINC, Parse Evercode Whole Transcriptome Kit v1.3.1 (Parse Biosciences, Seattle, WA), and 10X Chromium Single Cell 3’ Gene Expression Kit v3.1 (10X Genomics, Pleasanton, CA). These technologies employ distinct microfluidic and molecular barcoding strategies to capture the transcriptome of individual cells. To evaluate the comparative performance of these platforms, we analyzed separate cultures of DMSO-treated HepaRG cells. Due to differences in sample processing and data generation protocols between the Parse Evercode and 10X Chromium platforms, detailed descriptions of these protocols are provided in Supplementary Methods.

### Estimating the fraction of cells with responsive genes

To determine the fraction of treated cells showing significant upregulation of a gene compared to control cells, we used a statistical approach based on the negative binomial (NB) distribution, which is well-suited for modeling these data (Love et al., 2014). Given the expression levels across *n*_1_ treated and *n*_2_ control cells, we calculated the fraction (*f*_0_) of treated cells whose expression exceeded a threshold defined as 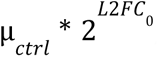, where µ_*ctrl*_ is the mean expression in control cells and *L*2*FC*_0_ is a predefined log₂ fold-change threshold (e.g., *L*2*FC*_0_ = 1 for a two-fold increase). To model gene expression in control cells, we estimated parameters for an NB distribution, which accounts for overdispersion in count data. Using the method-of-moments, we estimated the dispersion and probability parameters based on the mean and variance of control expression levels. Under the null hypothesis that treated cells follow the same distribution as controls, we calculated the probability that a cell’s expression exceeds the threshold using the NB survival function. We then used a binomial model to compute a p-value, representing the probability of observing at least as many upregulated cells by chance. This method provides both the fraction of treated cells for which the L2FC value of a gene exceeds the *L2FC* threshold and a statistical measure of confidence in the result. We refer to the fraction of cells for which the gene has L2FC>1 as FGT1+.

### Comparison with bulk differential expression

To compare chemical effects at the single-cell level with those expected in bulk measurements, we generated bulk (or "pseudo-bulk") transcriptomic profiles by aggregating single-cell data, as conventional bulk RNA-seq data were not available. The bulk profiles for TempO-LINC® samples were created by summing the raw gene counts from a subset of single cells exposed to the same chemical treatment. Specifically, three replicates were generated for each treatment by randomly sampling 33.33% of the single-cell transcriptomic profiles without replacement, and the summed gene counts were used to construct each pseudo-bulk profile. This approach consolidates transcriptomic data, reducing noise from single-cell variability while preserving dominant gene expression signals driven by chemical treatments. Differential expression analysis using the pseudo-bulk profiles was performed using DESeq2 (Love et al., 2014) to compare each treatment condition against the DMSO control. DESeq2 was used to calculate log2 fold change values and p-values for all genes, quantifying relative changes in gene expression. The p-values were adjusted for multiple comparisons using the false discovery rate (FDR) correction to obtain q-values (Benjamini & Hochberg, 1995). This analysis identified genes significantly upregulated or downregulated in the pseudo-bulk transcriptomic profiles as a response to chemical exposures, providing insights into affected biological pathways and processes. We refer to the differential gene expression est imated from bulk transcriptomic profiles as bL2FC.

### Relating transcripts to biological mechanisms and cell types

Understanding the transcriptional regulation of key pathways was critical for elucidating cellular responses to the chemicals tested in this study. Transcriptomic responses were related to specific biological mechanisms by focusing on key transcriptional targets within SRPs. The SRPs included UPR, DDR, OSR, HSR, HPX, and cell fate pathways such as APO, AUT, SEN, and peroxisome proliferator-activated receptor alpha (PPARα) signaling. Mechanisms were assigned based on key transcriptional targets identified through a comprehensive literature review. For the UPR, the transcription factors PERK (EIF2AK3) and IRE1A (ERN1) were focused upon. For the OSR, NRF2 (NFE2L2) targets were identified. The DDR was examined through the P53 (TP53) targets, and the HSR was studied by focusing on *HSF1*. Transcriptional targets for APO, AUT, CCA, and PPARα were identified based on existing literature. A detailed list of these targets for each mechanism is provided in Supplementary Table 1 (Table S1).

Identifying hepatocytes and cholangiocytes in SCTr data using transcripts relies on particular molecular markers. Albumin (ALB) serves as the gold standard marker for hepatocytes, with its liver-specific expression and abundance making it ideal for hepatocyte identification (Kmieć 2001). Hepatocyte nuclear factor 4-alpha (HNF4A) is a master transcription factor essential for hepatocyte differentiation and liver function (Walesky and Apte 2015). Additionally, apolipoprotein B (APOB), a key regulator of lipid metabolism and a major component of very-low-density lipoproteins (VLDL), is highly expressed in hepatocytes (Hussain, Strickland, and Bakillah 1999) and provides another robust marker for identifying this cell type. Arginase 1 (ARG1) shows high hepatocyte specificity as a critical urea cycle enzyme (Yan et al. 2010). For cholangiocytes, keratin 7 (KRT7) and keratin 19 (KRT19) represent the most reliable markers, with their consistent expression in biliary epithelium but absence in hepatocytes (Zhao et al. 2009). SOX9, a transcription factor crucial for bile duct morphogenesis, shows high specificity for cholangiocytes in the liver and is an excellent marker for biliary identification (Lin et al. 2022). CFTR and SCTR (secretin receptor) also demonstrate cholangiocyte specificity, with functional roles in bile secretion and modification (Venter et al. 2015). A detailed list of these and additional markers for each cell type is also provided in Table S1.

### Gene signatures of biological mechanisms

We also developed gene signatures for various biological mechanisms to complement individual gene markers. These signatures were constructed using a literature-mining approach based on the methodology outlined by Chambers et al. (B. A. Chambers et al. 2024). The process involved several key steps to identify relevant genes associated with specific biological mechanisms. First, we compiled a comprehensive list of all gene names and mechanisms (along with synonyms) and searched all PubMed abstracts for their co-occurrence. These mechanisms included OSR, UPR, HSR, HPX, DDR, AUT, SEN, and APO. Next, we quantified the strength of association between each gene and the mechanism by calculating normalized point-wise mutual information (nPMI) scores (Watford et al. 2018). The nPMI measures the statistical association between two terms, normalized to account for their frequencies. This approach allowed us to identify genes frequently mentioned in connection with a particular mechanism but also disproportionately associated with it compared to other genes. We applied a filtering step to refine the gene signatures, retaining only gene-term co-occurrences with an nPMI score greater than 0.2. This threshold excludes weak or spurious associations, ensuring the identification of meaningful connections between selected genes and biological mechanisms. Finally, we defined gene signatures of different lengths for each mechanism by selecting the top 50, 100, 150, and 200 genes based on their co-occurrence frequency and nPMI scores. These signatures were further validated against established genes (B Chambers and Shah 2021) for UPR, OSR, DDR, HSR, MSR, and HPX, ensuring consistency and relevance. The resulting signatures, linking 1,775 genes to SRPs, provide a robust tool for linking transcriptomic data to specific biological processes and are included in Supplementary Table 2 (Table S2).

### Connectivity scoring between gene signatures and transcriptomic profiles

Normalized single-cell transcriptomic (SCTr) profiles were standardized to facilitate treatment comparison and identify cellular responses relative to controls. Standardization was performed using the mean and standard deviation (SD) of the DMSO-treated control cells, generating a z-score (Z) for each gene in all chemical-treated cells. The z-score represents the differential expression of a gene, expressed as the number of SDs above or below the mean of the control, providing a quantitative measure of deviation from the baseline expression level. To capture the most responsive genes in each cell, the top 200 upregulated and 200 downregulated genes were selected based on their *Z* values, and this subset was denoted as *Z*^200^. The connectivity between *Z*^200^ and each SRP signature was calculated using the generalized Jaccard (gj) similarity metric. This metric measures the degree of overlap between gene signatures and profiles as a score (between -1 and 1) while accounting for differences in their relative expression levels, enabling the identification of pathways that are most closely associated with each cell’s transcriptional response. This approach allowed for the mapping of stress response activities for OSR, UPR, HSR, HPX, DDR, AUT, SEN, and APO at single-cell resolution. Further details about connectivity mapping are available in (Shah et al. 2022).

### Evaluating the significance of gene signature scores

The significance of chemical effects on SRPs measured by connectivity mapping and z-scores (henceforth referred to as “gene signature scores”) was assessed using two-way ANOVA to evaluate the impact of both chemical concentration and signature length. For each chemical-SRP combination, linear models were fitted with z-scores as the dependent variable and concentration and signature length as independent factors. While p-values (PR>F) from ANOVA indicated statistical significance of effects, due to the large sample size, effect sizes were quantified using eta-squared (η²) to determine practical significance. Eta-squared values, calculated as the ratio of factor sum of squares to total sum of squares, were interpreted as small (η² ≈ 0.01), medium (η² ≈ 0.06), or large (η² ≈ 0.14) effects (Richardson 2011). The direction and magnitude of effects were determined from the slope coefficients of the linear models, with positive slopes indicating activation and negative slopes indicating inhibition of the pathway with increasing concentration or signature length.

### Clustering and visualizing stress response states

Stress response activities for OSR, UPR, HSR, HPX, DDR, AUT, SEN, and APO were analyzed at single-cell resolution using *k*-means clustering to identify coherent patterns of stress response by reducing the dimensionality of the data. To determine the optimal number of clusters, multiple evaluation metrics, including silhouette scores (Rousseeuw 1987), Calinski-Harabasz index (Caliński and Harabasz 1974), and Davies-Bouldin index (D. L. Davies and D. W. Bouldin 1979), were computed and compared. These metrics provided quantitative assessments of cluster compactness and separation, ensuring robust identification of distinct groups of cells based on their stress response profiles. Cluster centroids, representing the average SRP activity of cells within each cluster, were further analyzed to explore relationships between clusters.

The relationships between clusters, SRPs, and connectivity scores were visualized using a custom heatmap, where clusters are represented as rows and SRPs as columns. SRP-cluster relationships were depicted as circles, with color saturation indicating the mean connectivity score and circle size representing the fraction of cells within each cluster that exceeded a connectivity score of 1 standard deviation above the mean for all cells in that SRP. Hierarchical agglomerative clustering was applied to the centroids (row-wise) and SRPs (column-wise) using the Euclidean distance metric and Ward’s linkage method, which minimizes variance within clusters during the merging process. The two-way hierarchical clustering of centroids and SRPs highlighted relationships between clusters and between SRPs. The heatmap circles effectively summarized the activity level and coherence of SRP responses within each cluster. This visualization provides a comprehensive overview of cellular SRP responses to chemical treatments, enabling the identification of patterns and trends in stress pathway activation across clusters.

### Analyzing putative cell state transitions

To analyze relationships between clusters, a putative cell state transition graph (CSTG) was constructed using two complementary approaches: (a) by iteratively connecting leaves within subtrees based on their proximity in the hierarchical clustering dendrogram, and (b) by linking each cluster to its two nearest neighbors based on centroid-to-centroid similarity. The resulting composite graph structure captured hierarchical relationships and direct transcriptional similarities among clusters. The CSTG graph was visualized using a force-directed layout, which arranges nodes based on simulated attractive and repulsive forces to enhance the interpretability of the cluster relationships. This dual-connectivity approach enabled a detailed mapping of cellular heterogeneity and putative stress response dynamics across the single-cell transcriptomic data.

### Implementation

All analyses on SCTr count data were performed in Python 3 using the Scanpy, scikit-learn, scipy, and seaborn libraries, along with custom packages (see SI1 for further information). The entire analysis can be accessed from GitHub (https://github.com/i-shah/sctr-i.git).

## Results

We present TempO-LINC data quality metrics and comparison with two main single-cell technologies using control samples to establish baseline performance. To assess chemical-induced responses, we quantified the proportion of cells expressing key marker genes across treatments. We applied single-cell visualization techniques to capture global transcriptional variation and identify treatment-associated effects. Single-cell gene signature analysis was used to evaluate stress response pathway activity, enabling the classification of cells based on adaptive response patterns. These data were clustered to define phenotypic subgroups representing putative cellular states. Finally, we constructed a cell state transition graph to organize these phenotypic groups to explore potential trajectories of cellular stress adaptation and progression.

### Data overview and quality

The TempO-LINC single-cell transcriptomics dataset comprised approximately 10 billion reads, with an average of approximately 150,000 reads per cell, derived from 48 samples. Each sample, on average, yielded 1,547 cells, contributing to a robust dataset totaling 33,742 cells that met our quality thresholds of over 500 genes and 10,000 (Supplementary Figure 1) read counts per cell. The unique molecular identifiers (UMIs) averaged 17,635 per cell across all samples in the dataset, with a high mapping rate of 85%, ensuring reliable gene expression data (Supplementary Figure S1B). The sequencing approach proved to be highly efficient, as we detected an average of 4,837 ± 114 genes per cell across the 22,533 probes used in the analysis (Supplementary Figure S1C). This high detection rate underscores the effectiveness of our sequencing approach. Notably, gene expression counts were uniformly distributed across all genes and cells within each treatment group (Table 2).

**Table 2.**
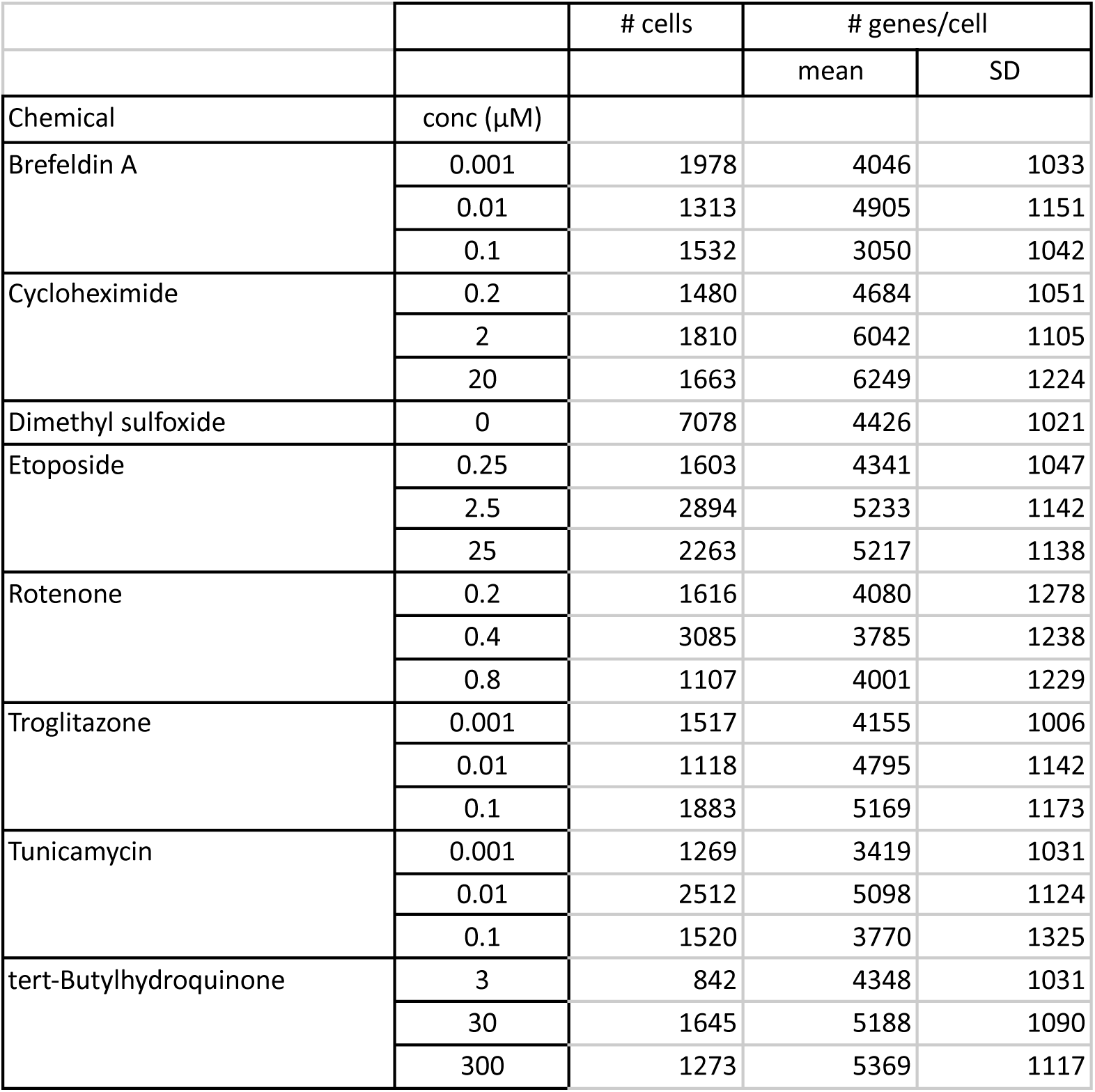
The number of cells and genes for each chemical treatment. The table shows the treatment chemicals and concentrations (conc) used in this study with the corresponding number of HepaRG cells (# cells), the mean number of genes per cell (# genes/cell) along with the standard deviation (SD).

### Platform comparison

A platform comparison was conducted to benchmark the performance of three single-cell transcriptomics (SCTr) technologies—BioSpyder TempO-LINC, Parse Evercode, and 10X Chromium —using control HepaRG cell samples. This comparison aimed to evaluate the relative strengths and limitations of each platform in terms of cell capture efficiency, library complexity, gene detection sensitivity, and overall data quality. The platform comparison was conducted on the negative control (DMSO-treated) samples using methods described in the Supplementary Methods. Gene expression metrics were determined following the normalization of transcript counts to an average of 5,000 transcripts per cell for each platform, ensuring comparability across the datasets generated by different technologies. This normalization step allowed a consistent baseline to evaluate key performance metrics such as gene detection rate and transcriptomic composition. The normalized data showing the distributions of genes and transcripts detected and distributions of the percentage of mitochondrial and ribosomal genes per cell are visualized in Supplementary Figure 2 (Figure S2). The analysis revealed that the TempO-LINC platform achieved a gene detection rate similar to Parse Evercode.

Furthermore, the transcriptomic composition analysis indicated that Parse Evercode had the lowest relative contribution of ribosomal RNA (rRNA), and TempO-LINC had the lowest mitochondrial RNA (mtRNA) transcripts (SF2A). Though this is useful, the rRNA and mtRNA can also be excluded from the data bioinformatically (SF2B). The TempO-LINC platform’s ability to exclude rRNA and mtRNA to prioritize biologically relevant and non-redundant transcripts with minimal interference from highly abundant housekeeping transcripts allowed us to focus on genes most relevant to the biological responses triggered by chemical stress, thereby enhancing the interpretability and impact of our findings. Additionally, the compatibility of the TempO-LINC assay with the TempO-Seq platform (v2.1), previously used for bulk cell high-throughput transcriptomics (HTTr) data (Bundy et al. 2024; Harrill et al. 2024). This compatibility ensures consistency across single-cell and bulk-cell analyses, enabling a comprehensive and integrated approach to studying chemical-induced responses. While not exhaustive, analyzing the same control HepaRG cell population with three different technologies directly evaluated platform-specific strengths and their suitability for single-cell transcriptomic applications.

### Single-cell responses to chemical treatments

To characterize chemical-induced effects at single-cell (SC) resolution, we quantified the fraction of cells exhibiting |L2FC| > 1 differential expression of all genes (FGT1+). This analysis revealed compound-specific response patterns and notable heterogeneity in pathway activation, highlighting the importance of SC resolution. As a baseline, we compared FGT1+ results with bulk differential expression (bL2FC), derived from aggregated single-cell counts. This comparison aimed to identify SC-specific responses potentially obscured in bulk analyses. By considering FGT1+ and bL2FC, we identified genes concordant between SC and bulk, and instances where subpopulation-specific effects, evident in SC data, might be diluted or masked in population averages. The number of significant genes (p<0.1) for each chemical treatment at different FGT1+ thresholds are shown in Table 3, and with details for specific genes provided in Supplementary Table 3 (Table S3). The effects of chemical treatments on genes are visualized using scatterplots of bL2FC vs FGT1+ in Figure 2, with highly induced genes annotated (The same scatterplots are visualized in Supplementary Figure S3, highlighting stress-response pathway-related genes). The following sections present illustrative results on chemical-induced gene expression.

**Figure 2.**
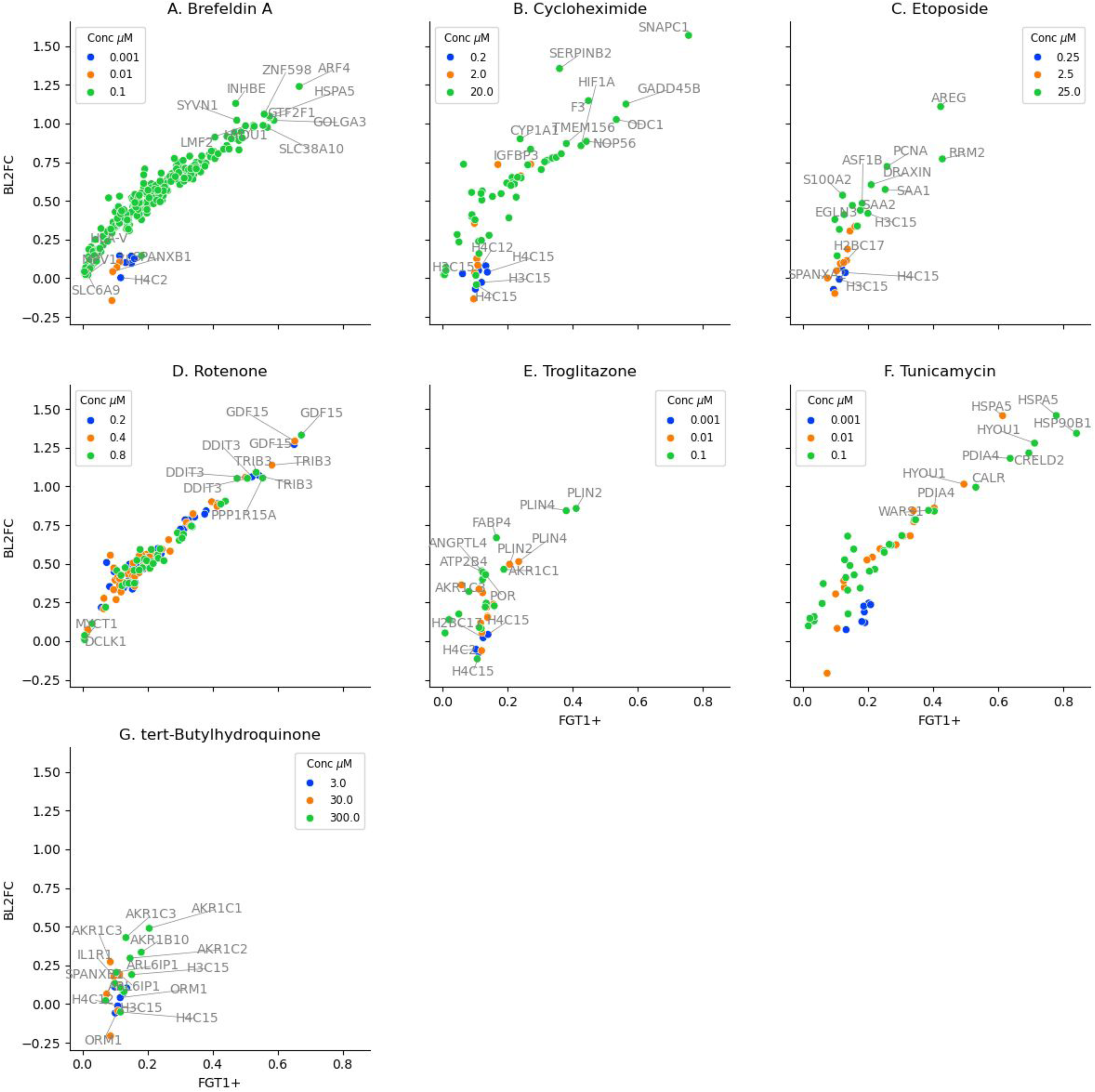
Chemical-induced transcriptional effects in single-cell and bulk data. The scatterplots show gene-level responses to each chemical as points, with the x-axis representing the fraction of responsive cells (FGT1) and the y-axis representing the bulk log₂ fold change (bL2FC). Each panel corresponds to a different chemical wih treatment concentrations represented by color. Only genes with statistically significant single-cell responses (p < 0.1) are shown. Illustrative examples of highly expressed genes and genes with heterogeneous expression (FGT1<0.2) are annotated in each scatterplot.

**Table 3.**
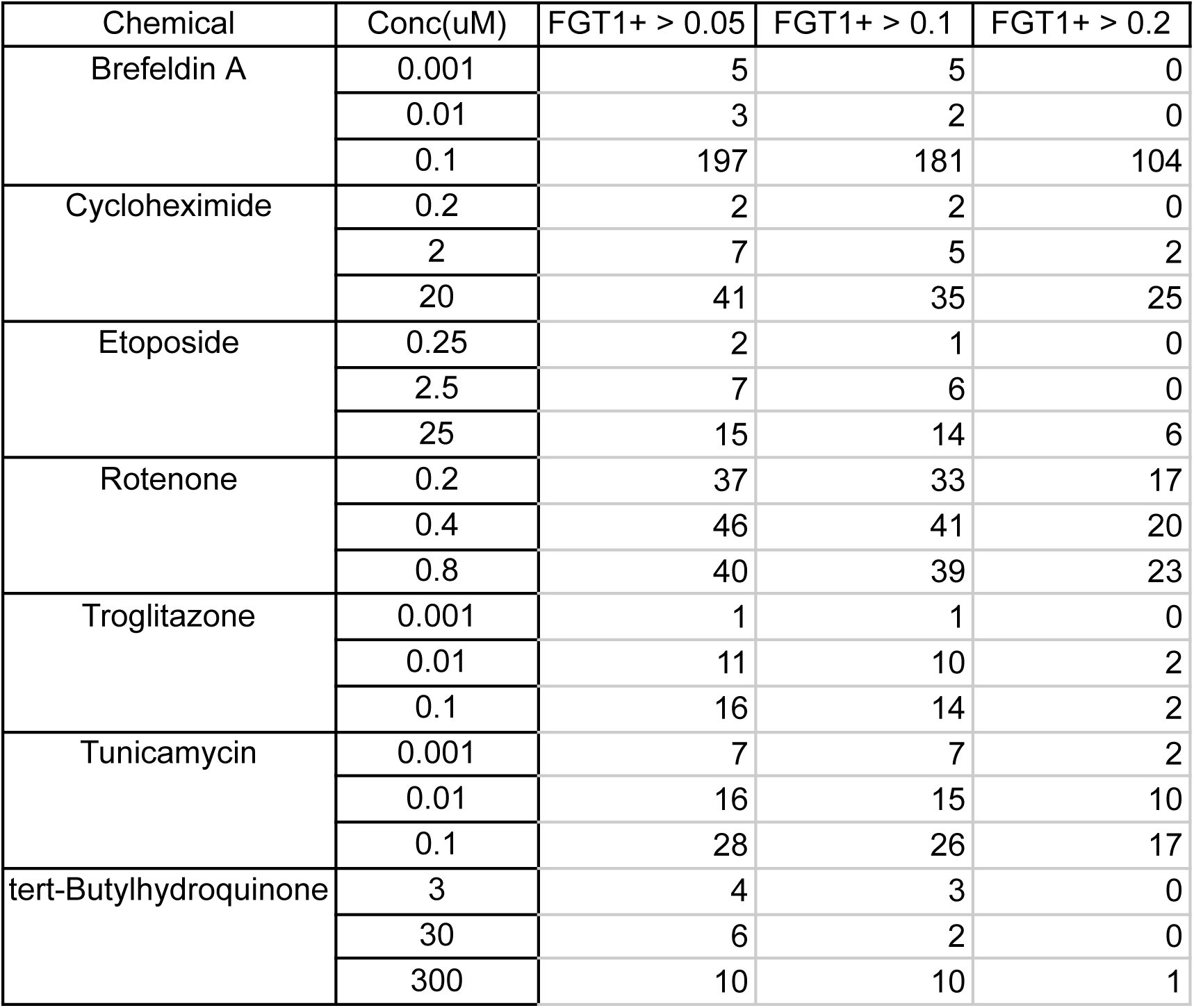
Summary of gene-level responses across chemical treatments and concentrations based on single-cell activity thresholds. This table reports the number of genes per chemical and concentration for which the fraction of responsive cells (FGT1) exceeded thresholds of 0.05, 0.1, and 0.2. Only genes with responses with p<0.1 in both single-cell and bulk data were included. FGT1 represents the proportion of treated cells in which the gene exhibited a log₂ fold change greater than 1, providing a quantitative measure of transcriptional activation at the single-cell level.

#### Brefeldin A effects

Brefeldin A, a fungal metabolite known to inhibit ER-to-Golgi protein transport (Sciaky et al. 1997), elicited a strong, concentration-dependent transcriptional response in HepaRG cells. At 0.1 µM, the highest concentration tested, 197 genes exhibited FGT1+ > 0.05, 181 exceeded FGT1+ > 0.1, and 104 exceeded FGT1+ > 0.2 (Table 3), indicating widespread and robust activation across the cell population. In contrast, at 0.001 and 0.01 µM, only 5 and 3 genes respectively showed FGT1+ > 0.05, with no genes exceeding 0.2, demonstrating minimal engagement of stress pathways at lower exposures.

Figure 2A shows the relationship between FGT1+ and bL2FC for genes significantly induced by Brefeldin A (p < 0.1). At 0.1 µM, the response was dominated by induction of genes associated with UPR, endoplasmic reticulum-associated degradation (ERAD), and proteostasis. HSPA5 (BiP), a central ER chaperone, was the most highly induced gene (FGT1+ = 0.58, bL2FC = 1.05), consistent with activation of the ATF6 and IRE1 branches of the UPR (Hetz 2012). GTF2F1 (FGT1+ = 0.58, bL2FC = 1.04), a general transcription factor, and GOLGA3 (FGT1+ = 0.59, bL2FC = 1.02), a Golgi scaffolding protein involved in ER-to-Golgi vesicle trafficking, were significantly upregulated. Their induction suggests coordinated adaptations to ER stress, including transcriptional reprogramming (Roeder 1996) and Golgi quality control during stress (Chang and Yang 2022).

Other highly expressed genes included SYVN1 (0.47, 1.02), an E3 ligase critical for ERAD (Lin et al. 2024), and LMF2 (0.50, 0.99), an ER membrane protein linked to lipid metabolism and ER homeostasis. The ER stress-associated chaperone HYOU1 (FGT1+ = 0.53, bL2FC = 0.99) and the amino acid transporter SLC38A10 (0.56, 0.99) were broadly induced, reflecting metabolic and proteostatic adaptation (Mann et al. 2024; Tripathi et al. 2019). ZNF598 (0.56, 1.06), involved in ribosome-associated quality control (Juszkiewicz et al. 2018), and PDIA4 (0.52, 0.89), an oxidoreductase supporting protein folding (Jiang et al. 2022), were also among the top responders. These findings point to coordinated activation of ER stress programs in the majority of cells.

SQSTM1 (p62), an autophagy adaptor, was upregulated in nearly half of the cells (FGT1+ = 0.49, bL2FC = 0.91), indicating induction of proteolytic clearance mechanisms, potentially through p62-mediated delivery of damaged proteins to lysosomes (Bjørkøy et al. 2005). Notably, a subset of cells exhibited selective expression of stress response transcription factors involved in apoptotic signaling. DDIT3 (CHOP) was expressed in 34% of cells (FGT1+ = 0.34, bL2FC = 0.64), and ATF4 in 30% (FGT1+ = 0.30,bL2FC=0.72), consistent with activation of the PERK-eIF2α-ATF4 branch of the UPR and progression toward pro-apoptotic signaling in prolonged ER stress (Ron and Walter 2007). INHBE, a stress-induced member of the TGF-β family, was also highly induced (FGT1+ = 0.47, bL2FC = 1.13), and it may contribute to paracrine signaling and cell fate regulation.

The concentration dependence of the response and the heterogeneity of gene induction underscore the value of single-cell resolution. While many key adaptive genes were broadly activated, pro-apoptotic and metabolic stress markers were limited to more minor subpopulations, revealing early divergence in cell state transitions that are not captured in bulk averages. These findings support the notion that Brefeldin A elicits both protective and potentially terminal cellular programs depending on the degree of stress and cell-intrinsic susceptibility.

#### Cycloheximide effects

Cycloheximide, a well-known inhibitor of eukaryotic translation elongation, elicited a concentration-dependent transcriptional response in HepaRG cells, consistent with cellular adaptations to global protein synthesis arrest (Schneider-Poetsch et al. 2010). At 0.2 µM, only two genes showed FGT1+ > 0.1, and none exceeded 0.2, whereas at 2.0 µM, five genes exceeded FGT1+ > 0.1 and two surpassed 0.2. At 20 µM, the highest concentration tested, the response was substantially broader, with 41 genes exceeding FGT1+ > 0.05, 35 above 0.1, and 25 above 0.2.

Figure 2B presents the correlation between FGT1+ and bL2FC for genes significantly upregulated by Cycloheximide (p < 0.1). Among the most strongly induced genes was ODC1 (FGT1+ = 0.53, bL2FC = 1.03), which encodes ornithine decarboxylase, the rate-limiting enzyme in polyamine biosynthesis. *ODC1* is tightly regulated at the translational and post-translational levels and is rapidly induced under stress conditions that affect protein synthesis. Its upregulation in response to Cycloheximide, a translation inhibitor, likely reflects compensatory transcriptional activation to maintain polyamine homeostasis and support cell survival under translational repression (Pegg 2006; 2009). *F3* (Tissue Factor) was also broadly induced (FGT1+ = 0.45, bL2FC = 1.15). As a key initiator of the coagulation cascade and a modulator of inflammatory signaling, elevated *F3* expression may reflect an early pro-inflammatory response or cellular damage signaling (Drake et al. 1989; Mackman 2004).

Several stress-responsive genes were induced across 30–45% of cells. These included NOP56, a ribonucleoprotein involved in rRNA processing (FGT1+ = 0.44) (Watkins et al. 2000), and HIF1A (FGT1+ = 0.43), a hypoxia-inducible transcription factor that may be stabilized by reduced protein turnover. SERPINB2 (FGT1+ = 0.36, bL2FC = 1.36), an inhibitor of plasminogen activators and a marker of cellular senescence and injury, was also upregulated, suggesting a shift toward cellular senescence or injury responses (Shaikh et al. 2024).

Other notable genes included GADD45A (FGT1+ = 0.31, bL2FC = 0.75), a DNA damage-inducible gene that mediates growth arrest and repair, and ATF4 (FGT1+ = 0.24, bL2FC = 0.65), a central mediator of the integrated stress response (Fulda et al. 2010). The induction of ATF4, in particular, reflects downstream activation of the eIF2α pathway triggered by translation inhibition (Harding et al. 1999). Additionally, IGFBP3, involved in growth factor signaling and apoptosis, and CYP1A1, a xenobiotic metabolism gene regulated by oxidative stress and aryl hydrocarbon receptor signaling, were highly expressed in ∼25% of cells.

These findings suggest that at high concentrations, Cycloheximide initiates a multi-layered response that integrates translation inhibition, metabolic stress, inflammatory signaling, and elements of DNA damage response. The heterogeneity of gene induction across single cells likely reflects differences in sensitivity to translational blockade, accumulation of misfolded proteins, and compensatory stress signaling.

#### Etoposide effects

Etoposide, a chemotherapeutic agent that induces DNA double-strand breaks via topoisomerase II inhibition <u>(Hande 1998)</u>, triggered a modest but mechanistically coherent transcriptional response in HepaRG cells. A dose-dependent trend was observed: at 0.25 μM, only 2 genes exceeded FGT1+ > 0.05 with none above 0.2; at 2.5 μM, 7 genes exceeded FGT1+ > 0.05 and 6 surpassed 0.1; while at the highest concentration (25 μM), 15 genes were induced above FGT1+ > 0.05, with 14 exceeding 0.1 and 6 above 0.2 (Table 3).

Figure 2C illustrates the relationship between single-cell responsiveness and bulk expression changes for Etoposide-induced genes (p < 0.1). At 25 μM, the top responding genes included RRM2 (FGT1+ = 0.43, bL2FC = 0.77), which encodes a subunit of ribonucleotide reductase essential for deoxyribonucleotide synthesis and DNA repair. Its induction reflects an increased nucleotide demand following DNA damage <u>(Giang et al. 2023)</u>. AREG (FGT1+ = 0.42, bL2FC = 1.11), a ligand of the EGFR pathway and transcriptional target of p53, suggests stress-induced regenerative or survival signaling <u>(Taira et al. 2014)</u>. Other moderately induced genes included PCNA (FGT1+ = 0.26, bL2FC = 0.72), a marker of DNA replication and repair <u>(Kowalska et al.</u> <u>2018)</u>, and ASF1B, a histone chaperone involved in chromatin remodeling during S-phase stress <u>(Groth et al. 2007)</u>. The acute-phase inflammatory mediators SAA1 and SAA2 were expressed in 18–25% of cells with moderate bulk upregulation, possibly reflecting paracrine signaling or cytokine activation downstream of genotoxic stress (Thorn et al. 2003).

Several genes were induced in more restricted subpopulations. For example, EGLN3 (FGT1+ = 0.15), a hypoxia-inducible gene implicated in apoptosis and stress adaptation, and S100A2 (FGT1+ = 0.12), involved in calcium signaling and modulating P53 activity (Mueller et al. 2005), showed low-frequency induction, suggesting cellular heterogeneity in damage resolution. These low-FGT1+ responses, while statistically significant, are unlikely to be captured in bulk profiling but may serve as early indicators of damage-associated cell states.

Altogether, Etoposide produced a focused transcriptional program involving DNA damage response, chromatin remodeling, and limited inflammatory signaling, primarily at high concentrations. The single-cell data highlight heterogeneity in response, with a minority of cells engaging stress or apoptotic pathways, consistent with p53-dependent transcriptional regulation (Vousden and Lu 2002). These observations reinforce the importance of resolving population substructure in evaluating the full spectrum of genotoxic responses.

#### Rotenone effects

Rotenone, a mitochondrial complex I inhibitor, produced a dose-dependent transcriptional response in HepaRG cells (Figure 2D), consistent with activation of oxidative and metabolic stress pathways <u>(Li et al. 2003)</u>. At 0.2 µM, 37 genes exceeded FGT1+ > 0.05, with 33 above 0.1 and 17 above 0.2. The response increased slightly at 0.4 µM (46 genes for FGT1+> 0.05; 41 genes for FGT1+>0.1; 20 genes for FGT1+> 0.2), and remained high at 0.8 µM with 40 genes above FGT1+ > 0.05, 39 for FGT1+> 0.1, and 23 genes for FGT1+> 0.2 (Table 3).

Figure 2D illustrates the significant effects of rotenone at a single-cell and bulk level (p<0.1). At a concentration of 0.8 µM, the most robustly induced genes included GDF15 (FGT1+ = 0.67, log2FC = 1.33), PPP1R15A (GADD34) (0.55, 1.05), TRIB3 (0.53, 1.09), and DDIT3 (CHOP) (0.51, 1.05). These genes are key effectors of the integrated stress response (ISR) and are regulated downstream of ATF4 following mitochondrial dysfunction or endoplasmic reticulum (ER) stress. Their strong induction across a large fraction of cells indicates a coordinated transcriptional response to proteotoxic and oxidative stress (Quirós et al. 2016; Harding et al. 2003).

PLIN2, which encodes a lipid droplet-associated protein involved in neutral lipid storage and mitochondrial metabolic adaptation, was induced in approximately 21% of cells (bL2FC = 0.59). This finding aligns with the metabolic reprogramming and lipid accumulation observed in hepatocytes under oxidative stress (Listenberger et al. 2003). HSPA5 (BiP), a chaperone that supports protein folding in the ER and is frequently induced under ER stress, was moderately upregulated (FGT1+ = 0.13, bL2FC = 0.37), indicating mild engagement of the unfolded protein response. While this level of induction was lower than seen with direct ER stressors like Brefeldin A, it supports the concept that mitochondrial ROS can secondarily perturb ER function (Zhao et al. 2002).

Overall, the SCTr data indicate that rotenone exposure activates a coordinated network of stress response pathways in a dose-dependent manner, with effects spanning the integrated stress ISR, oxidative stress, lipid metabolism, and mild engagement of UPR. At higher concentrations, a large fraction of the cell population exhibited strong induction of ISR effectors, reflecting widespread activation of ATF4-dependent transcriptional programs. In parallel, subsets of cells showed evidence of metabolic reprogramming and lipid accumulation, while others engaged proteostasis mechanisms. This is consistent with prior findings that mitochondrial complex I inhibition induces ROS production and downstream activation of stress and survival pathways (Sherer et al. 2003).

#### Troglitazone effects

Troglitazone, a thiazolidinedione and peroxisome proliferator-activated receptor gamma (PPARγ) agonist (Zhao et al. 2002), induced a modest but biologically relevant transcriptional response in HepaRG cells consistent with its known effects on lipid metabolism and hepatotoxicity (Figure 2E). Across the concentration range tested, the number of responsive genes increased from 1 gene (FGT1+ > 0.05) at 0.001 µM, to 11 at 0.01 µM, and 16 at 0.1 µM, with 14 genes exceeding FGT1+ > 0.1 and only 2 for FGT1+>0.2 at the highest dose.

At 0.1 µM, the top induced gene was AKR1C1 (FGT1+ = 0.19, bL2FC = 0.46), a member of the aldo-keto reductase family involved in steroid and xenobiotic metabolism. Its upregulation suggests early engagement of Phase I detoxification pathways (Penning and Drury 2007). FABP4 (FGT1+ = 0.17, bL2FC = 0.67), a fatty acid binding protein and canonical PPARγ target, was also significantly induced, supporting Troglitazone’s role in promoting lipid handling and adipogenic programs (Furuhashi & Hotamisligil, 2008). Other metabolic and lipid-associated genes included ANGPTL4 (FGT1+ = 0.12, bL2FC = 0.45), which modulates triglyceride clearance and is transcriptionally regulated by PPARγ (Biterova et al. 2018), and CD36 (FGT1+ = 0.02), a fatty acid transporter associated with lipid uptake and oxidative stress (Hua et al. 2015; Glatz et al. 2016). Their induction highlights a transcriptional profile reflective of PPARγ pathway activation and early shifts in lipid homeostasis.

POR (FGT1+ = 0.13, bL2FC = 0.43), encoding cytochrome P450 oxidoreductase, and AKR1C3 (FGT1+ = 0.12, bL2FC = 0.40) were also upregulated, pointing to enhanced capacity for xenobiotic metabolism, a hallmark of troglitazone’s hepatocellular effects (Masubuchi 2006). The calcium transporter gene ATP2B4 (FGT1+ = 0.12, bL2FC = 0.45) was induced in a small subset of cells, potentially reflecting early alterations in intracellular signaling or ionic stress (Giacomello et al. 2013).

Although gene induction levels were moderate and restricted mainly to subpopulations, these effects are consistent with known hepatotoxicity mechanisms of Troglitazone, including mitochondrial dysfunction, oxidative stress, and steatotic responses (Nanjan et al. 2018). The limited number of genes with FGT1+ > 0.2 suggests that while responses were specific, they were not widespread across the cell population.

#### Tunicamycin effects

Tunicamycin, a classic inducer of endoplasmic reticulum (ER) stress through inhibition of N-linked glycosylation (X. Chen et al. 2023), elicited a robust and concentration-dependent transcriptional response in HepaRG cells (Figure 2F). At 0.001 µM, only 7 genes showed FGT1+ > 0.05, with 2 for FGT1+>0.2. This response expanded at 0.01 µM (15 genes > 0.1, 10 > 0.2) and peaked at 0.1 µM with 28 genes for FGT1+>0.05, 26 for FGT1+> 0.1, and 17 for FGT1+> 0.2. These results are consistent with the broad activation of ER proteostasis and UPR pathways associated with tunicamycin exposure (Guha et al. 2017).

At 0.1 µM, the most prominently induced gene was CRELD2 (FGT1+ = 0.70, bL2FC = 1.22), a known downstream effector of ATF6 signaling that functions as an ER stress-inducible secretory protein (Oh-hashi et al. 2009). PDIA4 (FGT1+ = 0.64, bL2FC = 1.18), a protein disulfide isomerase involved in oxidative folding of nascent proteins, and CALR (calreticulin) (FGT1+ = 0.53, bL2FC = 1.00), a calcium-binding ER chaperone, were also among the top responders, consistent with widespread induction of chaperone-mediated protein folding (Jiang et al. 2022; Michalak et al. 2009).

The aminoacyl-tRNA synthetase gene WARS1 (FGT1+ = 0.39, bL2FC = 0.85) was also highly induced. While traditionally associated with translational regulation, WARS1 has been shown to respond to UPR activation and may play a role in translational control during stress adaptation (Park et al. 2008). Additionally, DNAJC3 (FGT1+ = 0.27, bL2FC = 0.62), a co-chaperone known to inhibit PERK signaling, was moderately induced, suggesting active engagement of feedback mechanisms to modulate the UPR (Fulda et al. 2010).

These findings point to strong activation of the canonical UPR branches, particularly ATF6 and PERK-eIF2α-ATF4, which regulate the expression of chaperones and folding enzymes to mitigate the accumulation of unfolded proteins in the ER (Ron and Walter 2007). The degree of cellular engagement, with more than half of the cells expressing key markers such as CRELD2 and PDIA4, suggests a homogeneous and well-coordinated stress response across the population.

#### tert-Butylhydroquinone (tBHQ)

tBHQ, a synthetic phenolic antioxidant widely used as a food preservative, is a well-characterized activator of the NRF2-mediated oxidative stress response (Imhoff and Hansen 2010). In HepaRG cells, tBHQ elicited a modest and concentration-dependent transcriptional response, with 4–10 genes exceeding FGT1+ > 0.05 across the tested dose range. At 300 µM, 10 genes surpassed FGT1+ > 0.1 and only one gene for FGT1+> 0.2, indicating a relatively limited engagement of stress pathways in the majority of the cell population.

Among the most responsive genes were several members of the aldo-keto reductase (AKR) superfamily, including AKR1C1 (FGT1+ = 0.21, bL2FC = 0.49), AKR1B10 (0.18, 0.33), AKR1C2 (0.15, 0.30), and AKR1C3 (0.13, 0.43). These enzymes play key roles in detoxifying reactive carbonyl species and are established transcriptional targets of NRF2 (Penning 2017). Their induction suggests that a subset of cells initiated an antioxidant defense program in response to tBHQ exposure.

Other induced genes included ARL6IP1 (FGT1+ = 0.10, bL2FC = 0.20), a regulator of ER membrane curvature and potentially ER stress responses, and SPANXB1 (0.10, 0.13), which may reflect chromatin remodeling or derepression of normally silenced loci in response to stress. Histone variants H3C15 and H4C15 were also modestly induced (FGT1+ = 0.15 and 0.12, respectively), though their biological relevance in the context of tBHQ exposure remains unclear.

Despite tBHQ’s established use as a model NRF2 activator, the restricted magnitude and frequency of gene induction suggest that only a limited proportion of HepaRG cells mounted a detectable oxidative stress response for the concentrations that we tested. Overall, these findings are consistent with selective NRF2 pathway activation at higher doses of tBHQ, as evidenced by AKR gene upregulation, but also highlight the heterogeneity of cell responses.

#### Summary of SC effects

Across all chemical treatments, single-cell transcriptomic analysis revealed both strong concordance with bulk gene expression and key distinctions that highlight the added value of single-cell resolution. As shown in Supplementary Figure S4, which plots all genes with p < 0.1 across treatments, there is a strong overall correlation between the fraction of responsive cells and bulk differential expression, with a Pearson correlation coefficient of 0.94. This indicates that, for the majority of genes, changes observed in population-averaged expression are mirrored by widespread induction across individual cells.

The single-cell analysis revealed three key themes in the transcriptional responses to chemical exposure. First, there was strong mechanistic specificity, with distinct genes mapping to known pathways such as ER stress (*HSPA5*, *PDIA4*), DNA damage (*GADD45A*, *RRM2*), oxidative stress (*AKR1C1*), and lipid metabolism (*PLIN2*, *FABP4*). Second, the responses were highly dependent on the chemical and dose, with some compounds like Brefeldin A and Tunicamycin inducing broad, uniform responses, while others, such as tBHQ and etoposide, elicited more heterogeneous effects limited to subpopulations. Single-cell data analysis revealed subtle patterns in multiple stress-related pathways. These transcriptional changes within specific cell subpopulations may serve as early indicators of cellular adaptation, compensation mechanisms, or progression towards terminal states.

### Visualizing single-cell transcriptomic data

We analyzed relationships between chemical treatments and stress responses by visualizing the similarity between the SCTr profiles using a two-dimensional t-SNE plot shown in Figure 3. Cells treated with DMSO exhibited a diffuse distribution across the t-SNE space, reflecting the inherent heterogeneity of differentiated HepaRG cultures (Figure 3B). Cells treated with 0.001 and 0.01 µM Brefeldin A overlapped with the DMSO-treated population, whereas cells exposed to 0.1 µM Brefeldin A formed a distinct distribution, indicating a concentration-dependent divergence in transcriptional response (Figure 3C). For cycloheximide, cells treated with 0.2 µM closely resembled the DMSO-treated population, while the 0.01 and 0.1 µM treatments produced a distinct distribution, indicating a transcriptional shift at higher concentrations (Figure 3D). In the case of etoposide, cells exposed to 0.25 and 2.5 µM were largely aligned with the DMSO distribution, whereas 25 µM treatment resulted in a clearly distinct cluster (Figure 3E). All concentrations of rotenone (0.001, 0.01, 0.1 µM) formed a distinct cluster, which partially overlapped with the distribution observed for 0.1 µM cycloheximide (Figure 3F).

**Figure 3.**
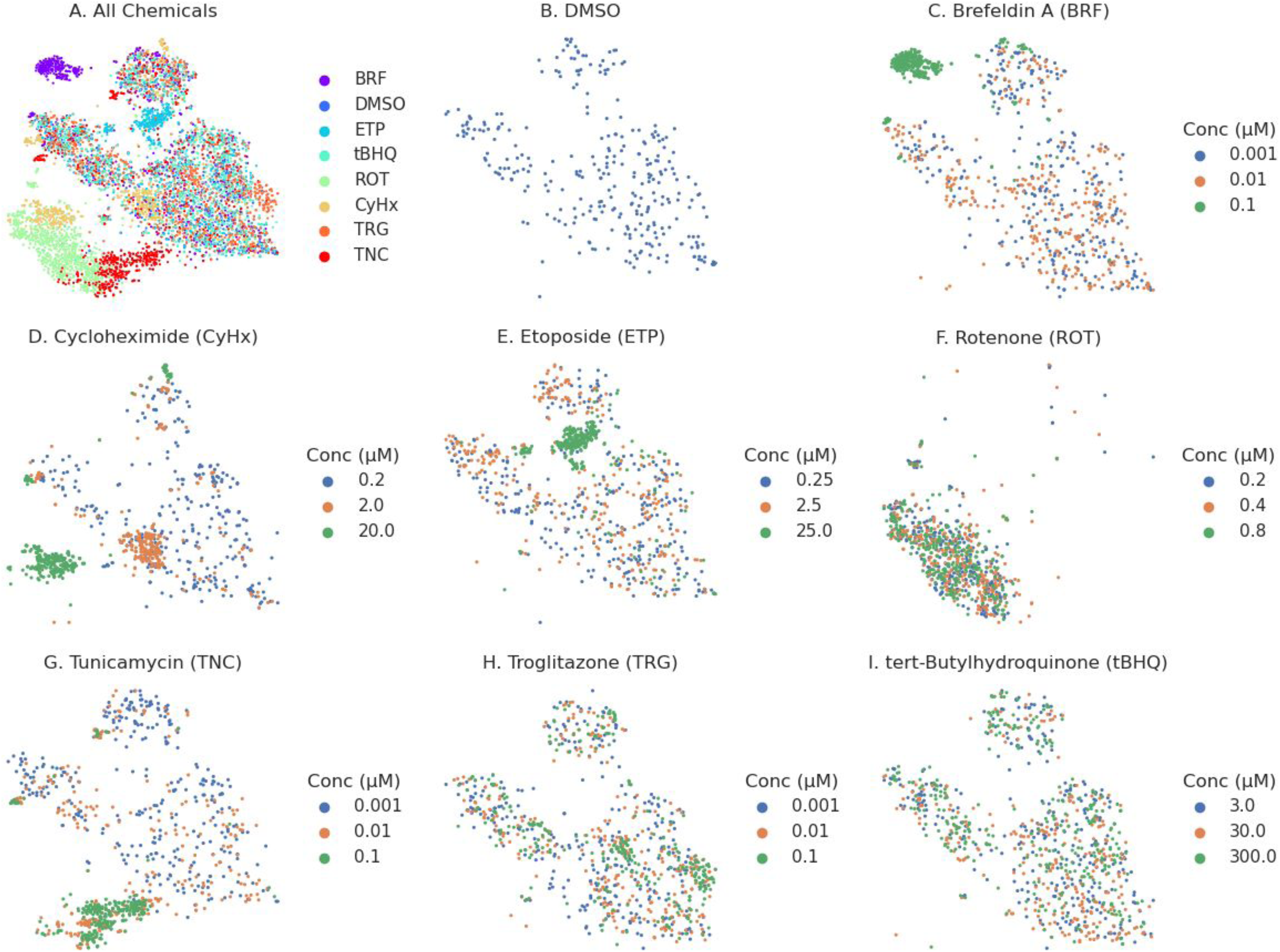
t-SNE plots of single-cell transcriptomic profiles. This figure shows t-SNE plots visualizing the transcriptomic profiles of individual cells in two-dimensional space. (A) Cells are color-coded by clusters and chemical treatments, showing how they group based on gene expression patterns and highlighting treatment-specific effects. (B) The expression levels of marker genes are depicted, with colors transitioning from deep blue (low or no expression) to red (high expression). These visualizations capture the diversity of cell states and reveal how treatments influence specific gene expression profiles.

For tunicamycin, cells treated with 0.001 µM closely resembled the DMSO group, while the 0.01 µM treatment resulted in a split distribution, partially overlapping with DMSO and partially forming a new cluster. Cells exposed to 0.1 µM tunicamycin aligned almost entirely with the 0.01 µM cluster (Figure 3G). Notably, the cluster formed by tunicamycin at 0.01 and 0.1 µM overlapped with the rotenone-associated cluster, suggesting some shared transcriptional responses.

Troglitazone-treated cells at all tested concentrations (0.001, 0.01, 0.1 µM) largely overlapped with DMSO-treated cells, although a focal enrichment of cells exposed to 0.1 µM was observed within a specific region of the t-SNE space (Figure 3H). Similarly, cells treated with tert-butylhydroquinone (tBHQ) at all concentrations predominantly co-localized with the DMSO population, indicating minimal transcriptional deviation under these conditions (Figure 3I).

The observed t-SNE distributions are consistent with the analyses of differentially expressed genes, where chemicals such as brefeldin A, tunicamycin, rotenone, etoposide, and cycloheximide showed strong induction of stress response pathways at higher concentrations. The distinct clusters for these treatments reflect widespread transcriptional activation in subpopulations of cells, particularly involving the UPR, HPX, and DDR. The partial overlap between rotenone and tunicamycin clusters corresponds with shared upregulation of UPR, suggesting convergent activation of proteotoxic and mitochondrial stress pathways. In contrast, the minimal shifts observed for tBHQ and troglitazone align with fewer differentially expressed genes and indicate weaker or more uniform responses. The spatial separation and overlaps among treatment groups in the t-SNE space provide a functional map of how different chemicals perturb adaptive stress pathways and highlight potential mechanistic similarities between treatments with shared transcriptional signatures.

Supplementary Figure S5 displays t-SNE plots of 15 genes with FGT1+<0.5 that mark distinct components of adaptive stress response pathways, revealing their heterogeneous expression across the cell population. Several genes are associated with the UPR (e.g., *HYOU1*, *SYVN1*, *PDIA4,* in Figures S5E, S5M, S5H, respectively), indicating activation of ER and proteostasis-related mechanisms induced by brefeldin A and tunicamycin. *DDIT3* (Figure S5B) and *INHBE* (Figure S5F) reflect pro-apoptotic and stress-induced signaling, while *RRM2* (Figure S5K) is linked to DNA damage repair and *NOP56* (Figure S5G) to ribosomal stress caused by etoposide and cycloheximide. Inflammatory mediators such as *F3* (Figure S5C) and *AREG* (Figure S5A) show some localized expression, pointing to damage-associated signaling due to multiple chemicals. *PLIN2* (Figure S5I) and *PLIN4* (Figure S5J) are involved in lipid metabolism and mitochondrial adaptation, and perturbed by troglitazone and rotetone. *WARS1* (Figure S5O) is an aminoacyl-tRNA synthetase upregulated under stress conditions. *HIF1A*, a key hypoxia-responsive factor, and *SERPINB2*, associated with senescence and injury responses due to rotenone and cycloheximide, further support the presence of multiple stress programs. These spatial expression patterns demonstrate how SCTr resolves cell-to-cell variability in SRP activation, capturing transcriptional diversity that is critical for understanding chemical-induced cell states.

The HepaRG cell cultures contain both hepatocytes and cholangiocytes, providing a potential model to examine cell-type-specific responses. To assess cell identity, we used a curated list of cell-type-specific markers (outlined in Methods and provided in Table S1) and visualized their expression on a t-SNE projection Supplementary Figure S6. The visualizations in Figure S6 allowed for an assessment of marker distribution across the cell population. Most markers did not form distinct clusters in the t-SNE space, indicating broad expression across cell types. However, two markers—APOB, ALB, and KRT19—showed notable expression patterns. ALB, a major plasma protein and hallmark of hepatocyte function, was highly expressed across a subset of DMSO-treated cells, consistent with a predominantly hepatocytic phenotype (Figure S6A). APOB, a key regulator of lipid transport primarily expressed in hepatocytes, showed a distribution pattern similar to ALB but was expressed across a slightly broader subset of DMSO-treated cells (Figure S6B). In contrast, KRT19, a marker of cholangiocytes and hepatic progenitor cells, exhibited a more restricted expression pattern that formed a partially complementary distribution to ALB and APOB, appearing in distinct subsets of DMSO-treated cells (Figure S6E). These observations suggest potential heterogeneous differentiation states within the HepaRG population, consistent with previous findings on its bipotential nature (Guillouzo et al., 2007).

### Cellular stress response analysis using gene signatures

A small set of gene biomarkers is likely insufficient to comprehensively capture the complexity of cellular responses, including adaptive stress mechanisms and phenotypic outcomes such as cell death. To better categorize cells by their phenotypic traits, we used gene signatures related to key stress response pathways (DDR, HPX, HSR, OSR, and UPR) and cell fates (AUT, SEN, and APO). We employed connectivity mapping to score single-cell transcriptomics profiles against these gene signatures, providing a simplified, lower-dimensional view of cellular responses (see Methods). Though multiple factors influence the analysis of transcriptomic data with gene signatures (Shah et al., 2023), we investigated two key variables: the impact of signature size and chemical concentration on stress response activities. The relevance of signature size stemmed from the possibility that larger gene signatures might capture broader pathway activation, whereas shorter signatures might offer more targeted insights into specific perturbations. Similarly, understanding the effect of concentration was critical for assessing dose-dependent shifts in pathway activation, albeit with a linear model, as only three treatment concentrations and the DMSO negative control were available for analysis. These factors (signature size and concentration dependence) were evaluated using two-way ANOVA, with the results provided in Supplementary Table 4.

Signature length showed statistically significant associations across multiple SRPs, but the effect sizes were consistently small (typical η² < 0.02). Notably, chemical effects on cell-level SRPs were reliably detected across varying signature lengths, particularly in the UPR, where chemicals such as Brefeldin A, Cycloheximide, and Troglitazone exhibited highly significant length-dependent effects (*p* < 1e-200) despite their small effect sizes. The distribution of scores for different signature lengths was very similar. Therefore, we use the results for a signature size of 200 for the remainder of the analysis. The standardized signature score (Z-score) distributions across chemical treatments and SRPs are shown in Figure 4. The DMSO treatments are shown as conc=0 for each treatment as a reference. Analysis of the relationship between chemical concentrations and SRPs revealed widespread effects, with most combinations showing significant concentration-dependence (*p* < 0.05). The strongest effects were observed for disruptors of protein synthesis and trafficking. Cycloheximide had a substantial impact on the UPR (η² = 0.148, slope = 0.053), consistent with its role in inhibiting protein synthesis. Brefeldin A strongly activated both AUT (η² = 0.103, slope = 10.19) and UPR (η² = 0.083, slope = 8.43), reflecting its disruption of protein trafficking. Tunicamycin significantly impacted UPR (η² = 0.093, slope = 10.68) and HSR (η² = 0.053, slope = -7.12), aligning with its role in inhibiting protein glycosylation. Troglitazone exhibited notable effects on OSR (slope = 1.59) and UPR (η² = 0.030, slope = 4.15). Etoposide showed strong activation of DDR (η² = 0.059, slope = 0.027), consistent with its mechanism as a topoisomerase inhibitor. Rotenone significantly affected APO (slope = 0.33) and AUT (η² = 0.036, slope = 0.77) pathways.

**Figure 4.**
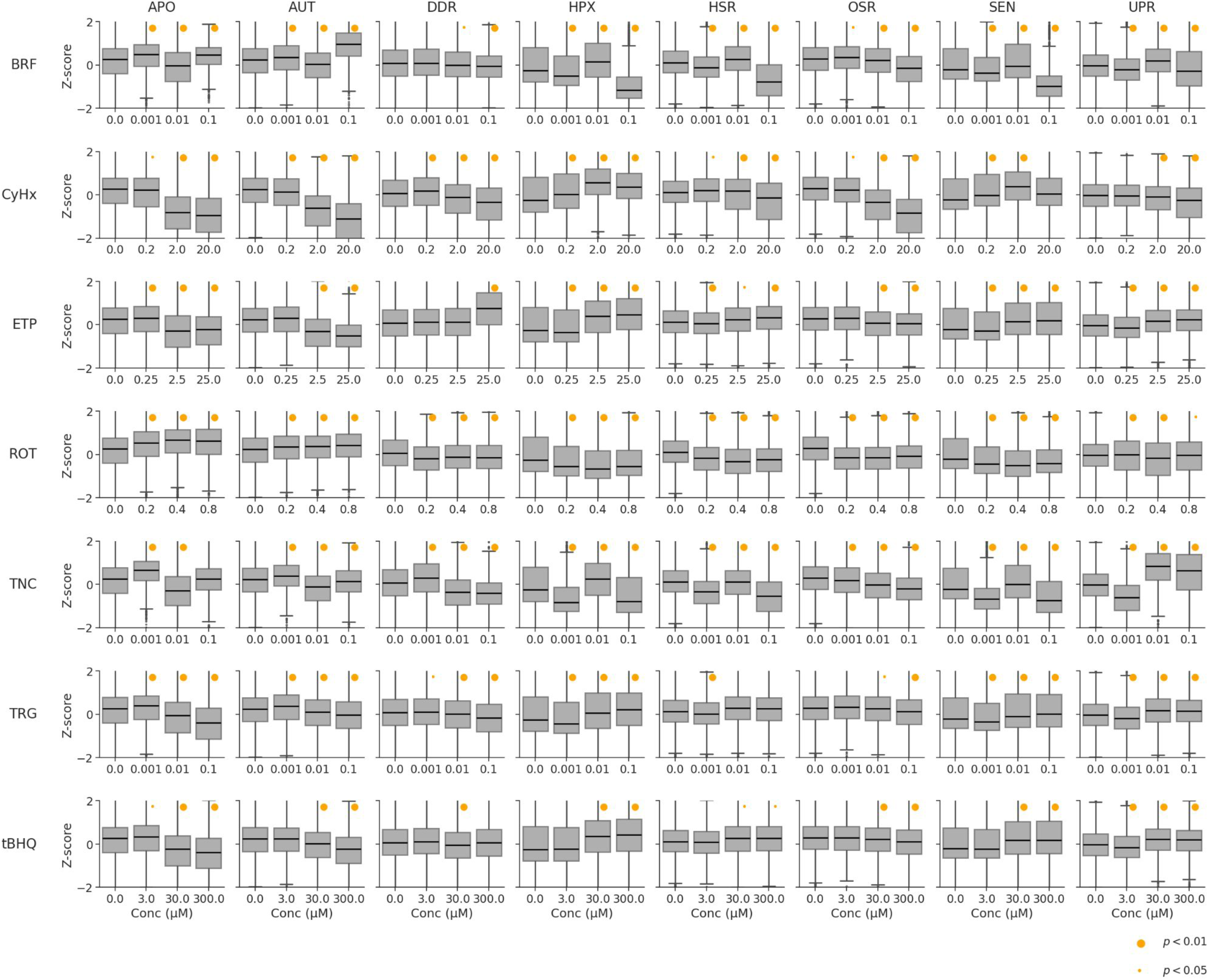
Connectivity score distributions for pathways across cells under different treatments This figure presents *z*-scores derived from generalized Jaccard scores, comparing single-cell transcriptomic profiles to pathway signatures. (A) Boxplots illustrate the distribution of *z*-scores for various chemicals (rows) across different concentrations, focusing on adaptive stress response pathways (columns), including apoptosis (APO), autophagy (AUT), DNA damage response (DDR), heat shock response (HSR), hypoxia response (HPX), oxidative stress response (OSR), senescence response (SEN), and unfolded protein response (UPR). (B) A heatmap summarizes the *z*-scores, capturing the activation levels of stress response pathways (SRPs) (columns) across the different chemical treatments (rows). These visualizations reveal the dose-dependent activation of specific pathways in response to chemical exposures.

The SRP gene signature analysis revealed significant secondary effects of chemicals beyond their primary SRP mechanisms, emphasizing the utility of comprehensive transcriptional profiling for uncovering broader cellular impacts. Cycloheximide, primarily recognized as a protein synthesis inhibitor, also demonstrated notable effects on DDR (*η²* = 0.071) and HPX (*η²* = 0.075), suggesting that its impact extends beyond translation inhibition to broader cellular stress responses, consistent with its known cytotoxic effects under prolonged exposure (Schneider-Poetsch et al., 2010). Similarly, Brefeldin A, while well-documented for disrupting protein trafficking, exhibited significant effects on APO signaling (*η²* = 0.082), indicating potential activation of cytotoxic pathways independent of its Golgi-disrupting activity, as supported by reports of its apoptotic effects in various cell types (Bladt et al. 2013).

Rotenone, typically associated with mitochondrial complex I inhibition, modulated AUT (*η²* = 0.036, slope = 0.77), reflecting the downstream impact of mitochondrial dysfunction on cellular stress pathways (Sherer et al., 2003). Tunicamycin, known for inhibiting N-glycosylation, also revealed substantial effects on HSR (*η²* = 0.053) and OSR (*η²* = 0.030), consistent with its ability to induce proteotoxic stress and oxidative damage through ER stress (Guha et al. 2017). Troglitazone demonstrated significant modulation of HSR (*η²* = 0.028, slope = 4.46) and UPR (*η²* = 0.030, slope = 4.15), suggesting a broader role in cellular stress responses potentially tied to its hepatotoxic effects (Fredriksson et al. 2014).

Gene signature analysis at the single-cell level matched chemicals to their known stress response mechanisms, demonstrating its utility in detecting biologically relevant perturbations across diverse SRPs. In some cases, responses were concentration-dependent, though effect sizes were generally small. The average single-cell signature activity scores at each chemical concentration aligned well with the known activities of the test chemicals, as reflected in the median distributions shown in Figure 4. However, this aggregate consistency masked the heterogeneity of cellular responses, as evidenced by the broad variability in single-cell activation scores, which can be seen in the tails of the distributions in Figure 4.

### Clustering cellular stress states into phenotypic groups

To further refine the analysis and capture highly responsive cells that might be overlooked in bulk transcriptomic data, we sought to identify distinct phenotypic groups within the SCTr dataset. We hypothesized that, despite differences in chemical treatment, individual cells would ultimately fall into a finite set of phenotypic states, reflecting the spectrum of adaptive stress and terminal responses. However, clustering the global SCTr data did not effectively distinguish between stress-adaptive and terminal response states (Figure 3), likely due to the high-dimensional complexity and variability of the full transcriptomic profiles. In contrast, the SRP signature scores provided a biologically meaningful lower-dimensional representation, effectively summarizing known chemical effects and stress response mechanisms. Therefore, we utilized the targeted 8-dimensional SRP signature score representation to capture the most relevant phenotypic states of cells. Clustering these SRP-based profiles allowed for the identification of distinct phenotypic groups, facilitating a more interpretable classification of cellular responses across different treatments.

To identify coherent cellular responses to different chemical treatments, we analyzed the 8-dimensional SRP signature scores at a single-cell level using k-means clustering. To determine the optimal number of clusters, we tested various partitions ranging from 10 to 100 clusters and evaluated them using multiple metrics, with the results visualized in Supplementary Information (SI13). The evaluation metrics, including silhouette scores and Calinski-Harabasz indices, consistently decreased as the number of clusters increased, suggesting the absence of a clear or perfect number of groups in the data. This trend indicates that the cellular states may be smoothly distributed across a continuum rather than forming discrete, well-separated groups. This inherent heterogeneity reflects the complex nature of stress responses, where cells may transition between states of adaptation and stress-induced death without discrete boundaries.

### Phenotypic groups

We selected *k* = 25 to balance granularity and interpretability in identifying phenotypic clusters, as shown in the rows of Figure 5A. The cell stress state clusters are visualized with circles whose color saturation reflects the mean signature score, and size represents the fraction of cells exceeding a score *Z* > 1 for a given SRP. Through hierarchical clustering, we manually classified the cell state clusters into five major phenotypic groups. This classification loosely defined groupings of related cellular states: Group I (clusters 8, 10, 16), representing a basal state with minimal SRP activation; Group II (clusters 1, 4, 5, 6, 11, 15, 18, 20, 21), showing selective APO and DDR activation; Group III (clusters 0, 2, 7, 9, 22, 24), characterized by coordinated activation of HSR, OSR, and UPR pathways with minimal APO and AUT; Group IV (clusters 3, 13, 12), displaying the highest intensity of HPX, SEN, and DDR responses; and Group V (clusters 14, 19, 23), distinguished by strong AUT and APO signatures. The hierarchical organization of cell states and their broader phenotypic grouping attempts to capture the relationships between different cellular states and provides a reduced dimension framework for understanding the spectrum of stress responses, from homeostatic conditions through increasing levels of adaptation to terminal stress states.

**Figure 5.**
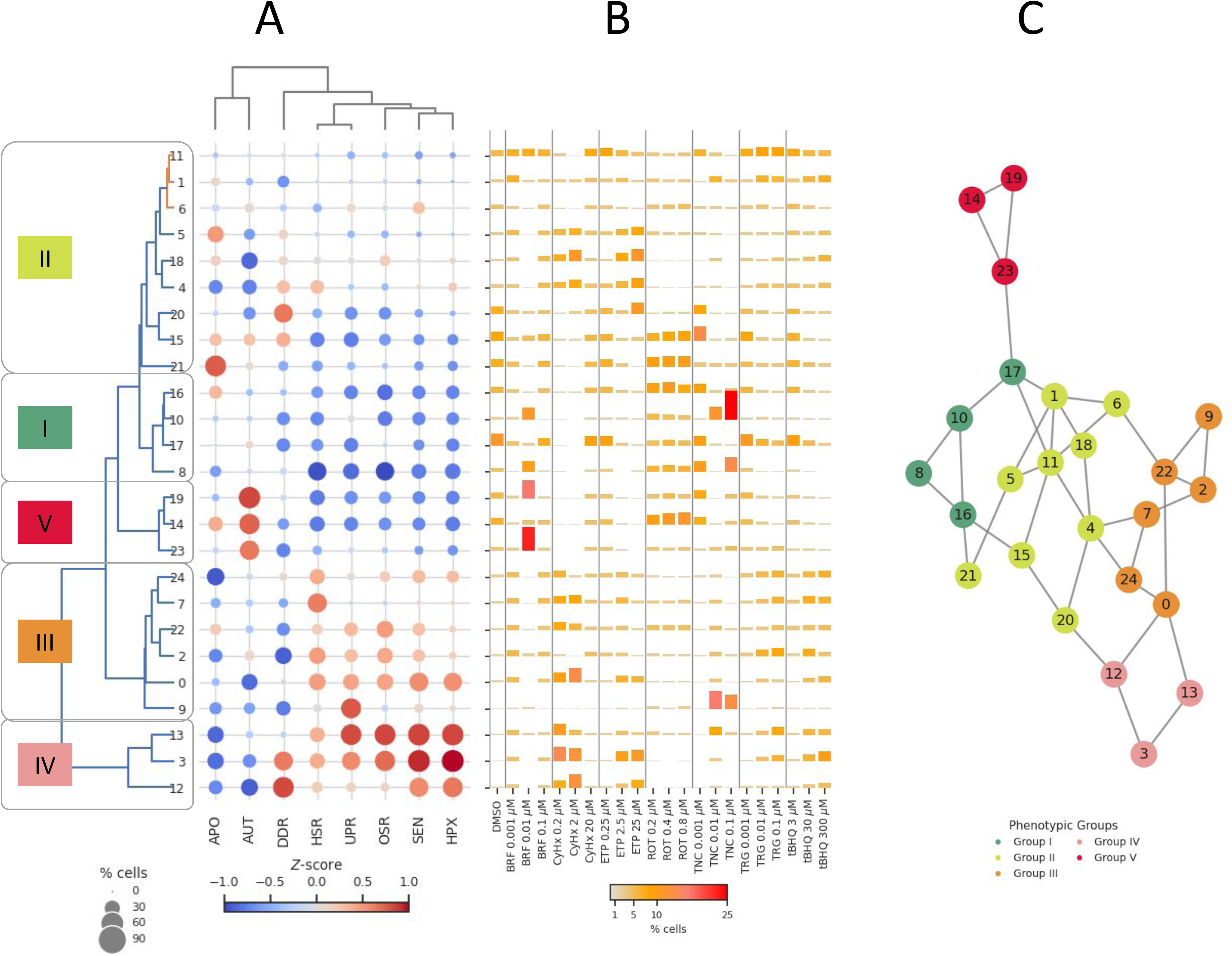
Stress response clusters and phenotypic groups. (A) A custom heatmap visualizes the relationships between clusters (rows), stress response pathways (SRPs) (columns), and connectivity scores. Circles represent SRP-cluster relationships, with color saturation indicating mean connectivity scores and size reflecting the fraction of cells in each cluster exceeding 1 standard deviation above the mean connectivity score for that SRP. (B) Barplots showing the distribution of the cell populations across clusters (rows) for each chemical treatment (columns).

### Heterogeneous cell stress responses

Chemical treatments exhibited diverse patterns of cellular stress responses across the phenotypic groups, reflecting substantial heterogeneity in how individual cells adapt to perturbation, as shown in Figure 5B. Some compounds drove cells predominantly toward specific stress states. For example, cycloheximide, a protein synthesis inhibitor, enriched cells in clusters 0 and 3 (14% and 12.4% respectively at 2.0 μM), which show high HSR/UPR activation consistent with its mechanism of proteostasis disruption. Similarly, tunicamycin, an ER stress inducer, demonstrated dose-dependent heterogeneity, with high concentrations (0.1 μM) driving cells predominantly to clusters 10 and 9 (27.6% and 13.4%, respectively), states characterized by UPR activation. In contrast, etoposide, a DNA-damaging agent, exhibited selective enrichment in clusters with high DDR activity, with substantial populations in clusters 18 and 3 (11.8% and 11.4% at 25 μM). Some cells from the same treatment activated targeted stress pathways, while others mounted broader adaptive responses or remained in basal states, indicating cell-to-cell variability in stress adaptation strategies.

### Visual evaluation of dose-response

A visual analysis of cellular responses revealed distinct patterns aligned with chemical stress mechanisms (Figure 5B). Cycloheximide, a protein synthesis inhibitor, demonstrated a biphasic response pattern. At intermediate concentrations (0.2-2.0 μM), cells progressively accumulated in states characterized by strong HSR and UPR activation, with up to 39% of cells occupying these stress states. However, this focused response dissipated at higher concentrations (20 μM) into a more uniform distribution, suggesting activation of alternative adaptation mechanisms or reaching a response threshold. Tunicamycin showed an apparent dose-dependent concentration of cells in UPR-specific states. While low concentrations (0.001 μM) resulted in a distributed response across various stress states, high concentrations (0.1 μM) drove a striking enrichment in UPR-dominant states, with approximately 41% of cells concentrating in these clusters. This pattern aligns with tunicamycin’s established role as an ER stress inducer.

Etoposide treatment revealed a progressive focusing of cellular responses toward DDR-specific states with increasing concentration. At its highest dose (25 μM), approximately 23% of cells concentrated in high DDR activity states, reflecting its mechanism as a DNA-damaging agent. In contrast, rotenone showed rapid achievement of a maximal response, maintaining a consistent enrichment of about 20% of cells in HPX/SEN states across all tested concentrations (0.2-0.8 μM), suggesting a threshold-type response to mitochondrial perturbation.

### Putative cell state transition graph

To explore relationships and transitions between cellular states, we constructed a cell state transition graph (CSTG) based on similarity in stress responses, following approaches established in single-cell trajectory inference studies (Trapnell et al. 2014; Saelens et al. 2018). The assumption that transcriptionally similar states represent more feasible transitions aligns with principles demonstrated in developmental biology research using single-cell data (H. Chen et al. 2019; Street et al. 2018). We used the 25 identified cell state clusters to build the CSTG (as detailed in Methods), incorporating both hierarchical relationships and direct transcriptional similarities to create a structured representation of potential cell state dynamics. The graph was overlaid with the five major phenotypic groups to contextualize transitions within broader adaptive stress response patterns (Figure 5C). By analyzing this CSTG, we can hypothesize putative cell state trajectories, providing insights into how cells may progress through different stress-adaptive and terminal response states. While comprehensive trajectory analysis exceeds this study’s scope, the CSTG establishes a foundation for future investigations into cell state transitions.

Chemical treatments induced highly heterogeneous and dose-dependent patterns of stress responses across cells, as evidenced by cluster distributions and phenotypic groupings. Some treatments, such as cycloheximide and tunicamycin, induced a dose-dependent change in the occupancy of cells in specific stress states aligned with their known mechanisms, including strong activation of HSR, UPR, and AUT pathways. In contrast, etoposide selectively enriched DDR-dominant states, while rotenone rapidly achieved a threshold response, consistently concentrating cells in HPX/SEN states. This variability highlights the distinct strategies cells employ to adapt to perturbations, with some treatments driving focused responses and others inducing broader distributions across multiple stress states. Overall, our findings show that while bulk analyses provide a helpful overview of overall stress activation, they obscure the nuanced, cell-specific dynamics, such as rare but highly responsive subpopulations or differential gene activation patterns.

## Discussion

This preliminary study demonstrates the feasibility and utility of SCTr in characterizing adaptive SRPs and cellular phenotypic heterogeneity in HepaRG cells following chemical exposure. By using the TempO-LINC® platform, we profiled the transcriptional responses of approximately 33,000 individual cells exposed to seven prototypical stress-inducing chemicals. These compounds were selected for their established ability to activate distinct SRPs, including the UPR, OSR, HSR, and DDR, allowing for broad coverage of toxicologically relevant mechanisms.

The use of the TempO-LINC platform proved effective for capturing biologically meaningful transcriptional responses in chemically perturbed HepaRG cells. The gene detection rate was consistent with other droplet-based SCTr platforms, and the targeted panel of probes used by TempO-LINC provided sufficient resolution to detect activation of known SRP markers and downstream fate-related genes such as those involved in apoptosis, autophagy, and cell cycle arrest. The current study also compared the performance of three platforms using the DMSO-treated controls, and our findings suggest that TempO-LINC’s gene detection rate was comparable to Parse Evercode and exceeded that of 10X Chromium.

The HepaRG model system offered several advantages for the analysis of stress responses in a human-relevant liver context. These bipotent progenitor cells differentiate into both hepatocyte- and cholangiocyte-like cells and retain many metabolic and transcriptional features of primary human hepatocytes, making them suitable for toxicogenomic studies (Guillouzo et al. 2007; Gerets et al. 2012). While their inherent heterogeneity is well documented, we were unable to detect a clearly distinct cholangiocyte subpopulation in our single-cell data. This may be due to incomplete or unstable differentiation of cholangiocyte-like cells under our culture conditions, selective loss of fragile or low-abundance cell types during fixation and dissociation, or limitations in marker resolution with the targeted panel used. Nonetheless, we observed transcriptional diversity across the HepaRG population, with some cells activating adaptive programs and others progressing toward terminal states in response to chemical stress.

Single-cell transcriptomic analysis at the individual gene level, conducted across diverse chemical treatments, showed significant agreement with bulk gene expression data. This high correlation suggests that population-averaged changes for most genes are indeed reflective of widespread induction across individual cells. Delving deeper, the single-cell data elucidated three key themes: first, a pronounced mechanistic specificity in transcriptional responses, with distinct genes clearly mapping to known stress pathways such as ER stress (e.g., HSPA5, PDIA4), DNA damage (e.g., GADD45A, RRM2), oxidative stress (e.g., AKR1C1), and lipid metabolism (e.g., PLIN2, FABP4). Second, we observed a strong dependence of response patterns on the specific chemical and its dose; for instance, lower doses of most compounds induced broad and uniform transcriptional changes, whereas higher doses elicited more heterogeneous effects restricted to distinct subpopulations. Finally, the single-cell approach uniquely uncovered fine-grained transcriptional heterogeneity within these subpopulations, often spanning multiple stress-related pathways, which could signify early adaptive or compensatory mechanisms, or transitions towards terminal cell states, thereby offering insights not apparent from bulk analysis alone.

Gene signature analysis revealed significant activation across multiple stress response pathways, and terminal cell outcomes such as autophagy and apoptosis. These findings underscore the broad engagement of adaptive and cell death programs following chemical exposure. Further partitioning of the single-cell data based on gene signature profiles identified five distinct transcriptional phenotypic groups. These ranged from a basal state with minimal SRP activation to progressively stressed states marked by pathway-specific engagement. Intermediate phenotypes displayed selective activation of individual SRPs, while the most perturbed groups showed co-activation of multiple stress responses and terminal outcome signatures. Chemicals like brefeldin A and tunicamycin prominently activated UPR markers, rotenone induced OSR consistent with mitochondrial inhibition (Sherer et al. 2003), and troglitazone elicited dose-dependent lipid metabolism changes alongside UPR and HSR, reflecting its known hepatotoxic effects (Fredriksson et al. 2014).

While established trajectory inference approaches such as partition-based graph abstraction (PAGA) can infer developmental or differentiation lineages based on transcriptional similarity, they are generally optimized for detecting gradual transcriptomic changes in well-defined biological processes (Wolf et al. 2019). In contrast, our goal was to investigate transitions between cell states defined by activation of specific adaptive SRPs, which may not follow smooth or lineage-like patterns. To this end, we developed a CSTG based on similarity in SRP pathway activity, using literature-derived gene signatures for UPR, OSR, HSR, and DDR. This approach allowed us to group cells based on mechanistically relevant stress responses rather than global transcriptomic profiles. While exploratory, the CSTG provides a structured framework to interpret how cells may transition from homeostatic to adaptive and ultimately to terminal states under increasing chemical stress, consistent with the concept of toxicological tipping points—thresholds beyond which sustained SRP activation may lead to irreversible cell outcomes such as apoptosis or autophagy (Shah et al. 2016; Hetz et al. 2020).

Although the proposed CSTG is not a definitive model of dynamic cell trajectories, it serves as a useful visual and analytical abstraction for interpreting high-dimensional SCTr data in the context of stress response biology. By grouping cells based on similarity in SRP pathway activity, the CSTG allows us to identify transcriptionally defined phenotypic states and infer plausible transitions between them. These connections provide a foundation for identifying potential tipping points, where cells may shift from adaptive stress responses to terminal fates, such as apoptosis or autophagy. This concept aligns with prior work on critical state transitions in cell fate decisions (Mojtahedi et al. 2016), which provides a concept of toxicological tipping points, where continued stress pushes a system beyond recovery. Observing these putative transitions at single-cell resolution underscores the value of SCTr in elucidating early molecular events and heterogeneous cell states that may underlie the mechanisms of chemical-induced toxicity.

Insights from the CSTG offer a nuanced perspective on how cells may transition from adaptive to terminal states in response to chemical stress. We observed that particular clusters, particularly those characterized by strong UPR, OSR, or DDR activation, appeared to serve as intermediate states that precede terminal phenotypes enriched for apoptotic or autophagic markers. This pattern supports the idea that sustained or excessive SRP activation can push cells beyond a recoverable threshold, aligning with mechanisms implicated in progressive liver pathologies such as steatohepatitis and hepatocellular carcinoma (Kanda et al. 2018; Spaan et al. 2019; Kouroumalis et al. 2021). By identifying phenotypic groups and their potential transitions, the CSTG provides a conceptual framework for distinguishing adaptive responses—where stress is managed and homeostasis preserved—from terminal outcomes, where damage becomes irreversible. This ability to map stress response trajectories at single-cell resolution could improve the mechanistic interpretation of in vitro assays, offering new opportunities to relate molecular signatures to phenotypic outcomes relevant to human liver toxicity.

It is also important to point out various limitations of the study presented here. For example, we observed distinct cellular responses even within the same treatment group, with some cells activating stress response pathways robustly while others maintained low expression of the same markers. This heterogeneity may reflect differences in cell cycle stage, intrinsic variability in stress sensitivity, or subpopulation diversity within the HepaRG model. Future studies incorporating lineage-defining markers or orthogonal modalities such as spatial or imaging-based profiling may help address this gap. Another consideration is the intertwined nature of stress response pathways and overlapping gene sets, which can complicate the interpretation of individual SRP activation patterns. For example, key transcriptional regulators such as ATF4 or DDIT3 participate in multiple stress pathways (Neill and Masson 2023), making assigning pathway activity exclusively to one biological process challenging. While gene signature–based scoring approaches help to summarize pathway activation (Chambers and Shah 2021), more refined models, such as network-based or probabilistic approaches, may be required to disentangle overlapping signals in future analyses.

The CSTG provided a valuable framework to visualize transitions between phenotypic states, but it is inherently limited by the static, single-time-point design of this experiment. Cellular states that arise early in the stress response or those that are transient may not be detectable at the 24-hour time point used here. Similarly, states that appear terminal in the CSTG may represent reversible phenotypes if sampled earlier or later. Incorporating additional time points or live-cell trajectory data would enable more accurate reconstruction of stress response dynamics and further clarify the temporal progression toward terminal cell fates.

Although this study profiled approximately 40,000 cells, it was designed as a preliminary investigation and included only a single HepaRG culture with one biological replicate per condition to minimize variability and focus on feasibility. This design limits our ability to assess inter-experimental variability and statistical robustness. Nonetheless, we observed strong and internally consistent patterns across chemical treatments, suggesting that key findings are biologically meaningful. Future studies incorporating additional replicates will be essential to strengthen confidence in pathway activity profiles and cell state transitions. Despite these limitations, this work highlights the utility of SCTr in resolving heterogeneous cellular stress responses and provides a foundation for deeper mechanistic exploration of chemically induced cell state dynamics.

While SCTr in monoculture systems such as HepaRG cells offers powerful resolution to identify stress pathway activation at the single-cell level, it inherently lacks the architectural and cellular complexity of the liver *in vivo* (Ishibashi et al. 2009). The liver is a highly structured organ composed of hepatocytes, cholangiocytes, Kupffer cells, hepatic stellate cells, liver sinusoidal endothelial cells, and a rich extracellular matrix, all contributing to immune surveillance, metabolic regulation, and repair processes (Trefts et al. 2017). These intercellular communications modulate stress responses in ways that are not captured in 2D monocultures. For instance, cytokine signaling from Kupffer cells can amplify hepatocyte responses to oxidative or ER stress (Imarisio et al. 2017), while stellate cells can initiate fibrosis in response to persistent damage (Puche et al. 2013). The absence of these paracrine and juxtacrine signaling pathways may influence the thresholds or dynamics of stress pathway activation observed *in vitro*, potentially underestimating toxicity or missing synergistic cell-cell effects. Our CSTG framework shows possible adaptive and terminal state transitions, but these interpretations are simplified by the monoculture system, which lacks critical immune and stromal factors found *in vivo*. However, this framework can be readily extended to more complex models, such as 3D liver spheroids (Ramaiahgari et al. 2017), or other microphysiological systems (Gough et al. 2021), where single-cell resolution will further enhance our understanding of tissue-level toxicological responses in a controlled and human-relevant setting. This direction holds promise for bridging the gap between molecular mechanisms and tissue-level pathologies.

From a translational perspective, SCTr offers several compelling advantages over traditional *in vivo* toxicology approaches. It enables high-throughput, cost-effective profiling of thousands of individual cells, revealing early molecular changes and heterogeneous responses that are often masked in bulk or animal-based studies. While animal studies remain essential for assessing whole-organism effects, they are time-intensive, expensive, and limited in mechanistic resolution (Kavlock et al. 2009). SCTr complements these methods by offering detailed insights into chemical-specific modes of action and potential tipping points in human-relevant systems. Platforms such as TempO-LINC, with targeted panels and simplified workflows, are lowering barriers to adoption by reducing per-sample costs and increasing throughput (Eastburn et al. 2024). As single-cell platforms continue to mature, combining SCTr with time-series sampling, imaging, and computational modeling will accelerate our ability to predict adverse outcomes from early stress responses (Shah and Wambaugh 2010; Sturla et al. 2014). As the field advances, the growing accessibility and scalability of SCTr will enable its broader application in predictive toxicology, complementing existing models and supporting a transition toward more human-relevant, mechanistically informed risk assessments.

Overall, the single-cell analysis presented here provides an early but compelling view of how HepaRG cells respond to chemical stressors at the level of individual phenotypes. By resolving the complexity of adaptive and terminal responses, SCTr holds promise for improving the interpretation of HTTr data by revealing how population-averaged signals emerge from diverse and dynamic cell states. While additional studies, including time-resolved designs, mechanistic perturbations, and 3D culture models, will be required to validate these findings, our results support the utility of SCTr for exploring stress response mechanisms and advancing the resolution and biological relevance of *in vitro* toxicity assessments.

## Conclusion

This preliminary study highlights the potential of single-cell transcriptomics to resolve adaptive and terminal stress responses in HepaRG cells exposed to chemical stressors. Using the TempO-LINC® platform, we identified distinct phenotypic states linked to major stress pathways, including UPR, OSR, HSR, and DDR. A putative cell state transition graph illustrated possible trajectories from homeostasis through adaptation to terminal outcomes. While additional validation is needed, these findings suggest that SCTr can uncover mechanistic transitions underlying toxicity and may complement high-throughput transcriptomics in advancing human-relevant NAMs.

## Acknowledgments

The authors would like to thank Drs. Jeffrey Dean and Joshua Harrill for technical review and constructive comments on this manuscript. Special thanks to Dr. Oswaldo Lozoya of the National Institute of Environmental Health Sciences for early discussions about this project, and to Jason Malphurs and Brian Pappas of the NIEHS Sequencing and Integrated Bioinformatics Cores for sequencing assistance and data handling.

## Disclaimer

The research described in this article has been reviewed by the US EPA and approved for publication. Approval does not signify that the contents necessarily reflect the views and the policies of the Agency. Mention of trade names or commercial products does not constitute endorsement or recommendation for use.

## Abbreviations

APO: Apoptosis
ATG5: Autophagy Related 5
AUT: Autophagy
BSA: Bovine Serum Albumin
CAT: Catalase
CSTG: Cell State Transition Graph
DDR: DNA Damage Response
DMSO: Dimethyl Sulfoxide
DSBs: Double-Strand Breaks
EBSS: Earle’s Balanced Salt Solution
EDTA: Ethylenediaminetetraacetic Acid
gj: Generalized Jaccard
HPX: Hypoxia
HSR: Heat Shock Response
HTTr: High-Throughput Transcriptomics
mtRNA: Mitochondrial RNA
NAMs: New Approach Methodologies
NCBI: National Center for Biotechnology Information
OSR: Oxidative Stress Response
PAGA: Partition-based graph abstraction
PBS: Phosphate-Buffered Saline
rRNA: Ribosomal RNA
SC: Single-Cell
SCTr: Single-Cell Transcriptomics
SD: Standard Deviation
SEN: Senescence
SI: Supplementary Information
SRA: Sequence Read Archive
SRPs: Adaptive Stress Response Pathways
t-SNE: T-distributed Stochastic Neighbor Embedding
tBHQ: Tert-Butylhydroquinone
UMI: Unique Molecular Index
UPR: Unfolded Protein Response
η²: Eta-squared

## Use of Generative AI

Grammarly’s writing assistant was used to enhance the readability of the prose by rephrasing sentences for grammatical correctness and clarity.

## Supplementary Information

### Supplementary Methods

Supplementary Table 1. Transcriptional (gene) markers of cellular pathways and cell types

Supplementary Table 2. Transcriptional (gene) signatures of cellular pathways

Supplementary Table 3. Comparison of single-cell vs bulk effects

Supplementary Table 4. ANOVA analysis of signature scores based on stress response gene signature and gene signature size

Supplementary Figure 1. Single-cell transcriptomics data quality

Supplementary Figure 2. Platform comparison

Supplementary Figure 3. Chemical-induced Gene Expression: Single-Cell vs Bulk Effects

Supplementary Figure 4. Relationship between the fraction of responsive cells and bulk differential expression

Supplementary Figure 5. Expression of key stress-related genes across single-cell populations

Supplementary Figure 6. Expression of key cell-type genes across single-cell populations

