## Supplementary Methods and Figures for "Decoding Cellular Stress States for Toxicology Using Single-Cell Transcriptomics"

SUPPLEMENTARY INFORMATION

Imran Shah[1*], David Gallegos[1^], Brian Robinette[1], Bryant A. Chambers[1#], Dennis J. Eastburn[2Δ], Douglas A. Bell[3±], Michelle R. Campbell[3], Suzanne N. Martos[3], Salvatore Camiolo[2], Kevin S. White[2], Nicole Martin[2], Gioele Montis[2], Joel McComb[2], Bruce Seligmann[2], Brian N. Chorley[1]

[1]Center for Computational Toxicology and Exposure, Office of Research and Development, U.S. Environmental Protection Agency, Research Triangle Park, North Carolina, USA.
[2]BioSpyder, Inc., Carlsbad, California, USA.

[3]Environmental Epigenomics and Disease Group, Immunity, Inflammation and Disease Laboratory, Intramural Research Program, National Institute of Environmental Health Sciences, National Institutes of Health, Research Triangle Park, NC, USA.

[^]Current Address: Takeda, Inc., Boston, MA, USA.

[#]Current Address: Inotiv, Inc., Morrisville, NC, USA.

[Δ]Current Address: Natera, Inc., San Mateo, CA, USA.

[±]Special Volunteer (retired), National Institute of Environmental Health Sciences - NIH

Table of Contents

**Supplementary Methods** 3

**Comparison with 10X Genomics and Parse Biosciences.** 3

10X Genomics Chromium Next GEM Single Cell 3’ preparation and sequencing 3

Parse Biosciences Single Cell Whole Transcriptome preparation and sequencing 4

Platform comparison 6

Implementation information 8

**Figure S1: Single-cell transcriptomics data quality** 10

**Figure S2: Platform comparison** 11

**Figure S3: Chemical-induced Gene Expression: Single-Cell vs Bulk Effects** 12

**Figure S4: Relationship between the fraction of responsive cells and bulk differential expression** 13

**Figure S5: Expression of key stress-related genes across single-cell populations** 14

**Figure S6: Expression of key cell-type genes across single-cell populations** 15

### Supplementary Methods

#### Comparison with 10X Genomics and Parse Biosciences.

Single-cell transcriptomic technologies provide unparalleled insights into the gene expression heterogeneity of complex cell populations. These tools have become invaluable for understanding cellular phenotypes and their responses to various stimuli. Among the most widely adopted platforms are the 10X Chromium Single Cell 3' Gene Expression Kit v3.1 (10X Genomics, Pleasanton, CA) and the more recent Parse Evercode Whole Transcriptome Kit v1.3.1 (Parse Biosciences, Seattle, WA). Both technologies employ distinct microfluidic and molecular barcoding strategies to capture the transcriptome of individual cells. To evaluate the comparative performance of these platforms, we analyzed control HepaRG cells using the 10X Chromium and Parse Evercode kits as described below.

##### *10X Genomics Chromium Next GEM Single Cell 3' preparation and sequencing*

An untargeted single-cell poly-A-tailed whole transcriptomic measurement using the Chromium Single Cell microfluidic platform was further utilized to assess the effect of DMSO exposure in HepaRG after 24 hrs. Single-cell preparations were as described in the "Cells dissociation & fixation" section in the main text, however, they were not fixed before library preparation. Briefly, single-cell HepaRG suspensions were barcoded and prepped for sequencing using the Chromium Next GEM Single Cell 3' Reagent Kit A (v3.1; 10X Genomics), following the manufacturer's instructions, with an input of 10,000 cells per sample. Then nanoliter-scaled Gel Beads-in-emulsion (GEMs) were partitioned to capture single cells as well as a master mix of Illumina TruSeq Read 1 primers, 10X barcodes, unique molecule identifiers (UMI), poly(dT) primers, and reverse transcription enzymes, followed by cell lysing and a cDNA synthesis reaction. After contents were released from the GEMs, pooled barcoded cDNA was purified by silane magnetic beads and amplified by PCR. After fragmentation, Illumina Read 2, P5, P7, and a sample index were added to the pooled amplified cDNA before paired-end sequencing on an Illumina NovaSeq 6000, using an S2 flow cell. Sample library prep traces were verified and quantified on a Bioanalyzer High Sensitivity DNA chip (Agilent).

Raw data generated using the 10X Chromium were processed following a standardized pipeline. Libraries were sequenced on an Illumina platform, producing paired-end reads with 28 bp for Read 1 (containing cell and molecular barcode information), 91 bp for Read 2 (cDNA sequences), and an 8 bp index read for sample identification. The resulting BCL files were converted to FASTQ format using the Cell Ranger mkfastq tool, which also demultiplexed libraries based on sample index barcodes. Reads were then aligned to the human reference genome (GRCh38) using the STAR aligner embedded within the Cell Ranger software suite. Transcript quantification was performed by extracting unique molecular identifiers (UMIs), filtering out low-quality reads, and collapsing UMIs to account for amplification artifacts, yielding a digital gene expression (DGE) matrix for each cell. Cells with fewer than 500 detected genes or excessively high mitochondrial RNA content (>20%) were flagged for potential exclusion in downstream analyses. The pipeline generated several key outputs, including a filtered feature-barcode matrix of high-quality data, a raw matrix for all detected barcodes and genes, and quality control metrics summarizing sequencing performance, such as median genes per cell and sequencing saturation. Filtered data were exported in Matrix Market format for downstream analyses. FASTQ files for these samples are available on the NCBI Gene Expression Omnibus (GEO) under accession # XXXXX.

##### *Parse Biosciences Single Cell Whole Transcriptome preparation and sequencing*

We also utilized a commercially available, single-cell whole transcriptomic “split-pool” approach using the Parse Biosciences Single Cell Whole Transcriptome kit (v1.3.1, Parse Biosciences, Seattle, WA). Like the other protocols, 24 hr rotenone exposure was performed in HepaRG using two doses (0.2 and 0.4 µM). To ensure enough cells were prepared for the protocol, 6-well plates instead of 24-well plates were used for culture, seeding at approximately $3.5X10^3$ cells/$cm^2$. Single-cell preparations were as described in the “Cells dissociation & fixation” section in the main text, however, they were fixed using the Parse Cell Fixation kit (v1.3.0) before library preparation, following the manufacturer’s instructions. After dissociation, fewer than 4 million cells were used for the fixation protocol to avoid increased doublet rates (averaged ~$9X10^5$ cells per sample at 93% viability). Cells were spun at 200 xg for 10 min at 4°C, and pellets were resuspended in kit Cell Buffer + 0.5% BSA. Cells were filtered through a 40 µm strainer and incubated in 4°C kit fixation solution after a 3X mix by pipetting and incubated on ice for 10 min. 4°C kit. Permeabilization solution was then added, mixed 3X by pipetting, and incubated 3 min on ice. Kit neutralization buffer was then added, the tube was inverted once to mix, and fixed cells were pelleted at 200xg for 10 min at 4°C. Pellets were then suspended to obtain a minimum concentration of 600 cells/µl, cells were then passed through a 40 µm strainer, and finally, DMSO was added slowly over 3 steps, with a 3X flick mix and 1 min incubation with each aliquot. Cells were then counted and then slow frozen to -80°C storage until prepped for sequencing.

Library preparation was performed with the Parse Biosciences Single Cell Whole Transcriptome kit (v1.3.1), following the manufacturer's instructions. Briefly, approximately $2X10^3$ cells were loaded per well, after thawing the fixed samples in a 37°C heat block. Samples were then barcoded in four "split, then pool" rounds. In round 1, cells were reverse transcribed, and barcodes were added in situ using primers. Samples were then pooled and redistributed, and another round (2) of barcodes was added by ligation. Again, samples were pooled and redistributed, and a third round of barcodes was added using an in situ round of ligation. After each pooling, cells were spun in a swing-bucket centrifuge for 10 min at 200 xg at 4°C, where indicated. Finally, a sublibrary barcoding was distributed using eight aliquots of the round 3 pool, adding 8 Illumina indices after lysing the aliquots. No more than $1.3X10^4$ cells were added to each sublibrary based on counts performed before the final barcoding. Afterward, barcoded cDNA was bead-purified, template-switched, amplified, and SPRI bead clean-up was performed. Sublibraries were then quality checked on a Bioanalyzer and quantified. Subsequently, 100 ng of each sublibrary amplified cDNA was enzyme fragmented, end-repaired, and A-tails were added. Fragmented sublibraries were then double-sided SPRI bead selected, indexing adapters were ligated, SPRI beads cleaned, and then amplified using Illumina indices. After a post-amplification double-sided SPRI bead clean-up, sequencing libraries were quality checked on a Bioanalyzer and quantified High Sensitivity DNA chip (Agilent). Sequencing was performed on the 8 sublibraries using the paired-end protocol on an Illumina NovaSeq 6000 with an S4 flow cell.

Raw data generated using the Parse Biosciences Single Cell Whole Transcriptome Kit v1.3.1 (Parse Biosciences, Seattle, WA) were processed following the Parse Biosciences pipeline, optimized for split-pool combinatorial barcoding data. Libraries were sequenced on an Illumina platform, producing paired-end reads with 150 bp for both Read 1 and Read 2, where Read 1 contained the combinatorial barcodes and Read 2 captured the cDNA sequence. The sequencing data were demultiplexed to generate FASTQ files for downstream processing. Raw FASTQ files were processed using the Parse Biosciences pipeline, which first identified and extracted cell-specific barcodes from Read 1 to assign reads to individual cells. Reads were then aligned to the human reference genome (GRCh38) using a splice-aware aligner to map gene expression accurately. Unique molecular identifiers (UMIs) were extracted, and PCR duplicates were removed by collapsing identical UMIs associated with the same gene. This process generated a digital gene expression (DGE) matrix representing the counts of unique transcripts per gene per cell. Quality control filtering was performed to remove low-quality cells, such as those with fewer than 500 detected genes or excessively high mitochondrial RNA content (>20%). The pipeline output included a filtered DGE matrix, gene and cell-level quality control metrics summary, and sequencing performance statistics. The resulting DGE matrix was exported in a standard format for downstream analyses in single-cell analysis tools such as Scanpy or Seurat, where normalization, dimensionality reduction, clustering, and visualization were conducted to investigate cellular heterogeneity and gene expression profiles. This processing protocol ensured the generation of high-quality, reproducible single-cell transcriptomic data using the Parse Biosciences split-pool barcoding approach. Raw sequencing data are available on the NCBI Gene Expression Omnibus (GEO) under accession # XXXXX.

##### *Platform comparison*

Platform comparison between BioSpyder TempO-LINC, Parse Evercode, and 10X Chromium was conducted using cells treated with DMSO based on gene, transcript detection rates, and the percentage of ribosomal and mitochondrial genes per cell. Raw count matrices were generated for each platform downstream of the data processing to generate raw counts. Raw counts were exclusively used for the analysis, with no normalization or scaling applied to the datasets. Platform-specific count matrices were used to create individual Seurat objects for downstream data processing. Prior to merging, the Seurat objects were filtered based on metadata annotations to retain only cells from DMSO-control-treated conditions. Cells with fewer than 5,000 UMIs were excluded from each dataset. To ensure consistent benchmarking, each platform-specific count matrix was then downsampled to a uniform depth of 5,000 UMIs using the `SampleUMI` command (`max UMI = 5000`). The downsampled datasets were annotated by platform and merged into a single Seurat object using the `merge` function. Gene quantification for mitochondrial, ribosomal, and hemoglobin transcripts was performed using the `PercentageFeatureSet` function, with the classifiers `'^MT-'` for mitochondrial genes, `'^RP[SL]'` for ribosomal genes, and `'^HB[^(P)]'` for hemoglobin genes. After initial comparisons, mitochondrial and ribosomal gene transcripts were removed from the dataset using string-based classification. Genes were identified using the `counts[which(grepl("^RP[SL]", rownames(counts))),]` command and removed from the downsampled object using the `subset` command. Following this filtering step, comparison metrics were re-generated using the revised downsampled dataset without mitochondrial and ribosomal transcripts. All analyses were conducted using R version 4.3.0 and Seurat version 5.0.1, ensuring consistency in the platform-specific and merged dataset processing workflows.

#### Implementation information

All analyses were conducted in Python 3 using a combination of libraries and custom scripts to facilitate data processing, visualization, and modeling. Single-cell transcriptomic analysis was performed using Scanpy (Wolf et al., 2018), a scalable toolkit for preprocessing, clustering, and visualization of high-dimensional single-cell data. Connectivity mapping analysis was conducted using the Generalized Connectivity Toolkit (gecco) package (Shah et al., 2022), which provides robust methods for evaluating relationships between gene signatures and stress response pathways (SRPs).

Data visualization utilized seaborn (Waskom, 2021) for statistical graphics, along with matplotlib (Hunter, 2007) and networkx (Hagberg et al., 2008) for hierarchical clustering and force-directed graph representations of cluster relationships. Statistical analyses, including regression models and significance testing, were implemented with SciPy (Virtanen et al., 2020), while machine learning algorithms and clustering methods, such as *k*-means clustering, were performed using scikit-learn (Pedregosa et al., 2011). For handling large-scale data, Dask (Rocklin, 2015) was employed to enhance parallel processing and computational efficiency.

This comprehensive computational framework combined specialized libraries with the gecco package to ensure rigorous and reproducible analysis of connectivity scores and cellular heterogeneity. The code is available from the authors upon request.

Wolf, F. A., et al. (2018). Scanpy: large-scale single-cell gene expression data analysis. *Genome Biology*, 19, 15.

Waskom, M. (2021). Seaborn: statistical data visualization. *Journal of Open Source Software*, 6(60), 3021.

Hunter, J. D. (2007). Matplotlib: A 2D graphics environment. *Computing in Science & Engineering*, 9(3), 90–95.

Hagberg, A. A., et al. (2008). Exploring network structure, dynamics, and function using NetworkX. *Proceedings of the 7th Python in Science Conference (SciPy2008)*, 11–15.

Virtanen, P., et al. (2020). SciPy 1.0: Fundamental algorithms for scientific computing in Python. *Nature Methods*, 17(3), 261–272.

Pedregosa, F., et al. (2011). Scikit-learn: Machine Learning in Python. *Journal of Machine Learning Research*, 12, 2825–2830.

Rocklin, M. (2015). Dask: Parallel computation with blocked algorithms and task scheduling. *Proceedings of the 14th Python in Science Conference (SciPy2015)*, 126–132.

Shah, Imran, et al. (2022). Navigating Transcriptomic Connectivity Mapping Workflows to Link Chemicals with Bioactivities. Chemical Research in Toxicology 35 (11): 1929–49.

#### Figure S1: Single-cell transcriptomics data quality

SIA. Knee plot

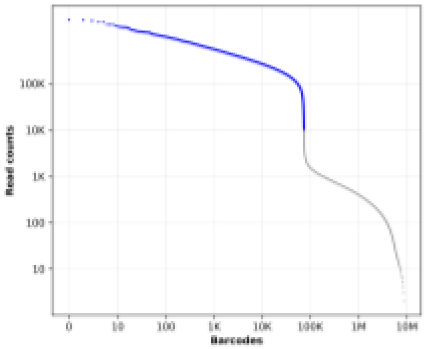


| Human Hepatocytes: 10,000 reads per cell threshold | | | | |
|---|---|---|---|---|
| Number of Reads | Number of UMI | Mapping Rate | Avg. # Cells/Sample | # Detected Genes |
| 148,642 ± 6706.7 | 17,635 ± 734.8 | 84.7 ± 0.6 % | 1547 ± 75.6 | 4837 ± 114.1 |

S1B. Read-depth, Mapped Reads and UMIs

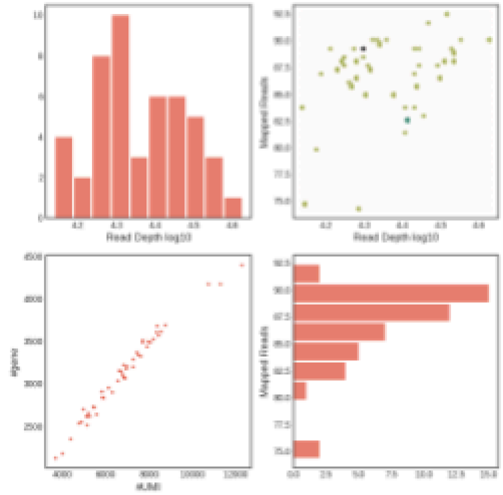


S1C. Number of Genes vs Reads Per cell

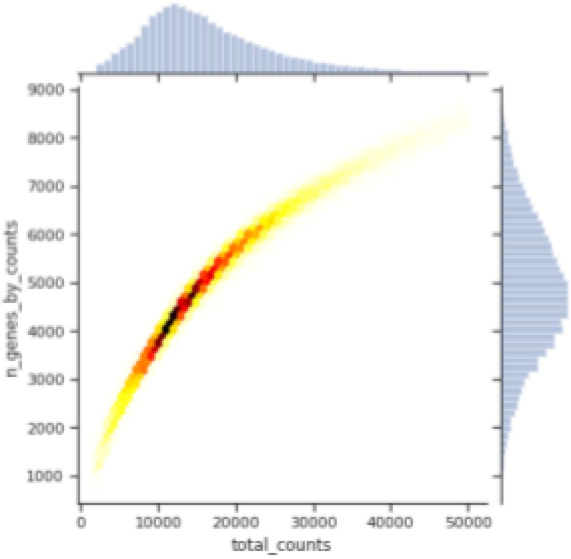


Evaluation of the quality of the single-cell transcriptomic data obtained using the TempO-LINC platform. Panel S1A illustrates a knee plot used to determine a threshold for cell calling, identifying a 10,000 reads per cell cut-off. Panel S1B shows metrics related to sequencing depth, including read-depth, the number of mapped reads, and Unique Molecular Identifiers (UMIs). Panel S1C depicts the relationship between the number of genes detected per cell and the number of reads per cell.

#### Figure S2: Platform comparison

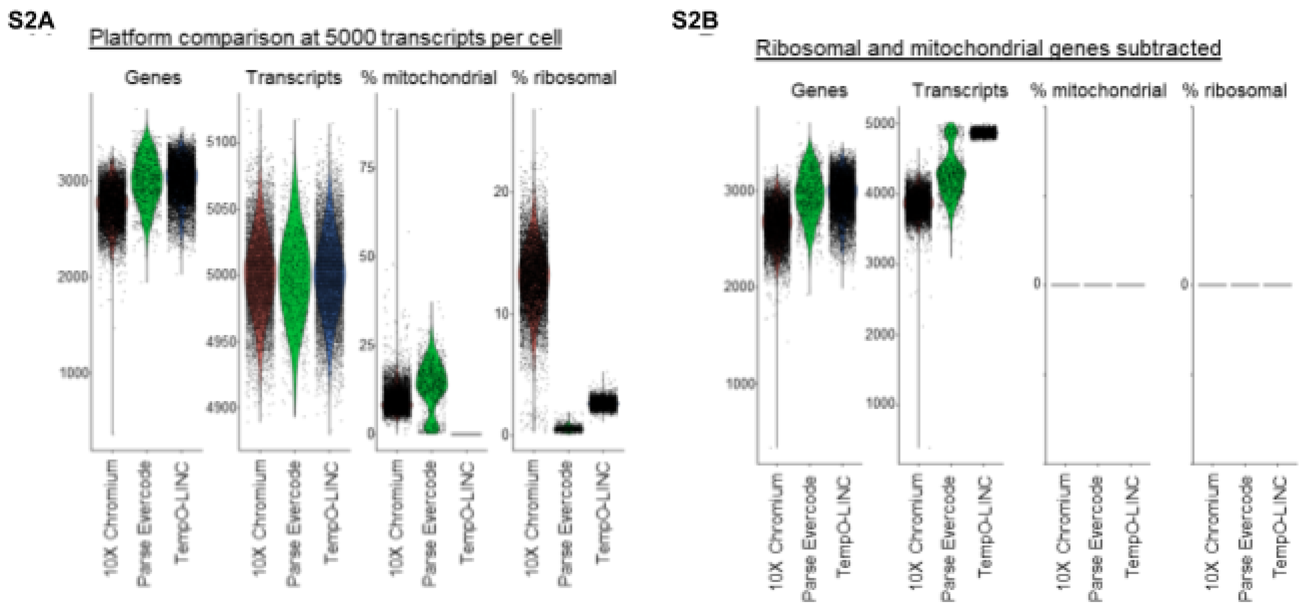


Performance comparison of three single-cell transcriptomics platforms: BioSpyder TempO-LINC, Parse Evercode (Whole Transcriptome Kit v1.3.1), and 10X Chromium Single Cell (3' Gene Expression Kit v3.1). Figure S2A shows the distributions of the number of genes per cell, number of transcripts per cell, percentage of mitochondrial genes, and percentage of ribosomal genes, all normalized to 5,000 transcripts per cell. Figure S2B presents the same distributions after subtracting mitochondrial and ribosomal genes from the analysis, allowing for a more focused comparison of gene and transcript detection.

#### Figure S3: Chemical-induced Gene Expression: Single-Cell vs Bulk Effects

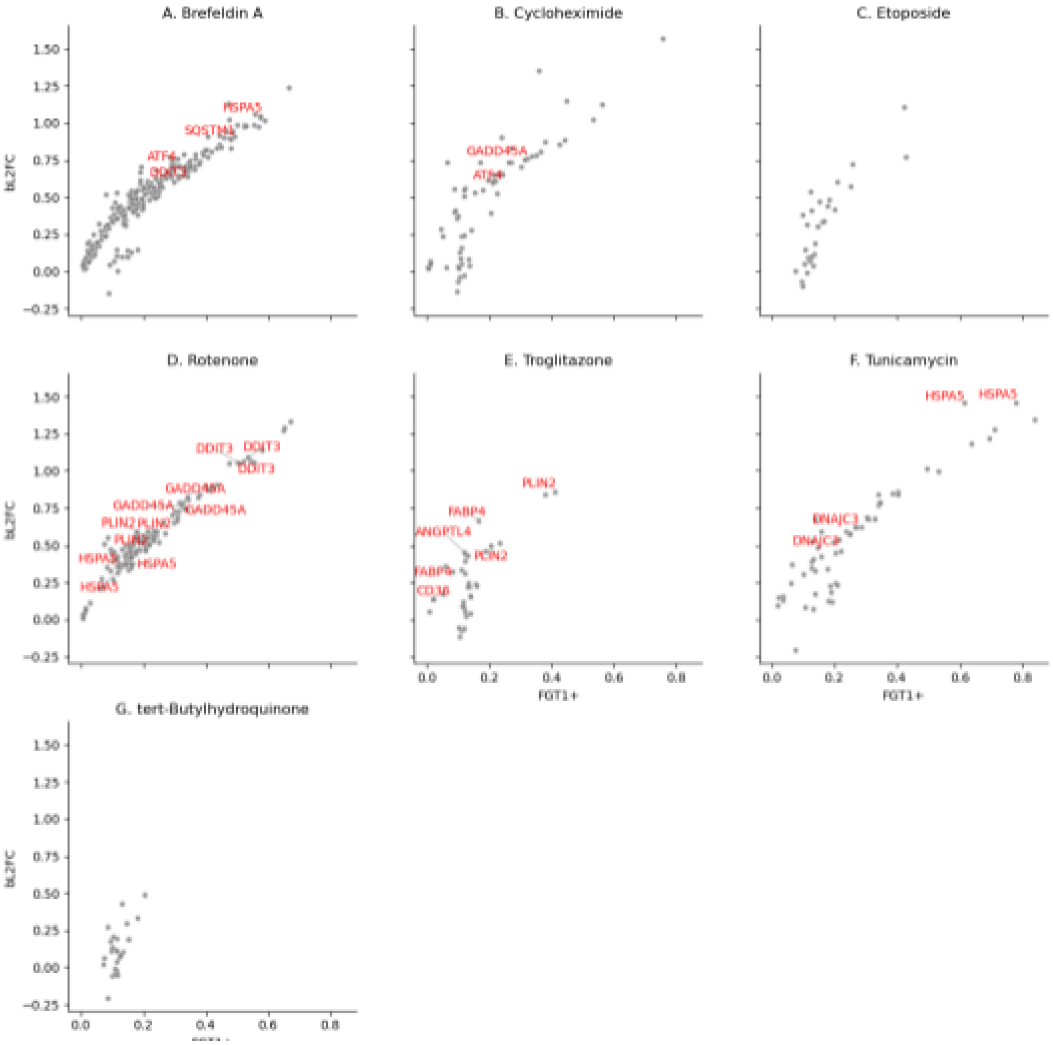


The scatterplots show gene-level responses to each chemical, with the x-axis representing the fraction of responsive cells (FCT1+) and the y-axis representing the bulk $\log_2$ fold change (bL2FC). Each panel corresponds to a different chemical treatment. Only genes with statistically significant single-cell responses ($p < 0.1$) are shown. Genes annotated as belonging to curated Stress Response Pathways (SRPs)—including UPR, OSR, HSR, DDR, APO, AUT, CCA, and PPARA—are labeled in red. All other genes are shown in grey. Annotated genes illustrate mechanistic relevance and pathway-specific transcriptional activation.

#### Figure S4: Relationship between the fraction of responsive cells and bulk differential expression

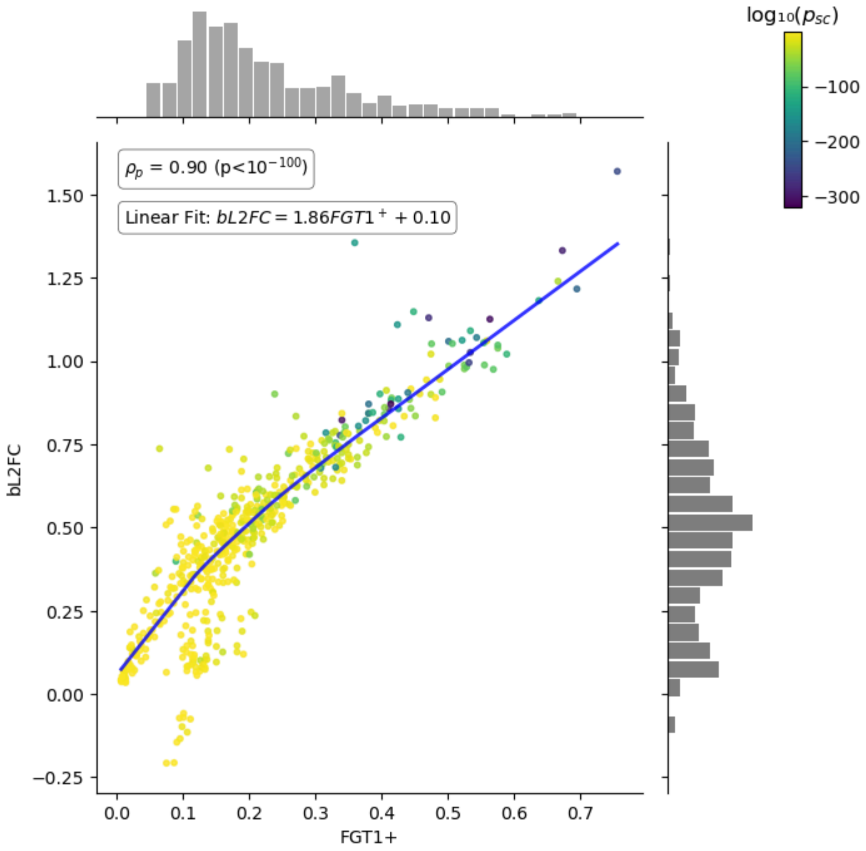


This scatterplot shows each gene's single-cell response (x-axis: FCT1+, the fraction of treated cells with $\log_2$ fold change > 1) against its bulk-level $\log_2$ fold change (y-axis: bL2FC). Only genes with statistically significant effects in both single-cell and bulk measurements (sc_p < 0.1 and blk_p < 0.1) are shown. Points are color-coded by the $\log_{10}$-transformed p-value from the single-cell test ($\log_{10}$(p□$_c$)) using a viridis colormap. A linear regression line is overlaid with the equation *bL2FC = 1.53·FCT1⁺ + 0.07*, and a LOWESS (locally weighted scatterplot smoothing) curve is shown in blue. The Pearson correlation coefficient (ρ□) was 0.94 ($p < 10^{-100}$), indicating a strong linear relationship. Marginal histograms (30 bins) show the distribution of values along each axis, with counts plotted on a linear scale and labeled as "n". A colorbar is positioned in the lower right to illustrate the $\log_{10}$-scaled single-cell p-values.

#### Figure S5: Expression of key stress-related genes across single-cell populations

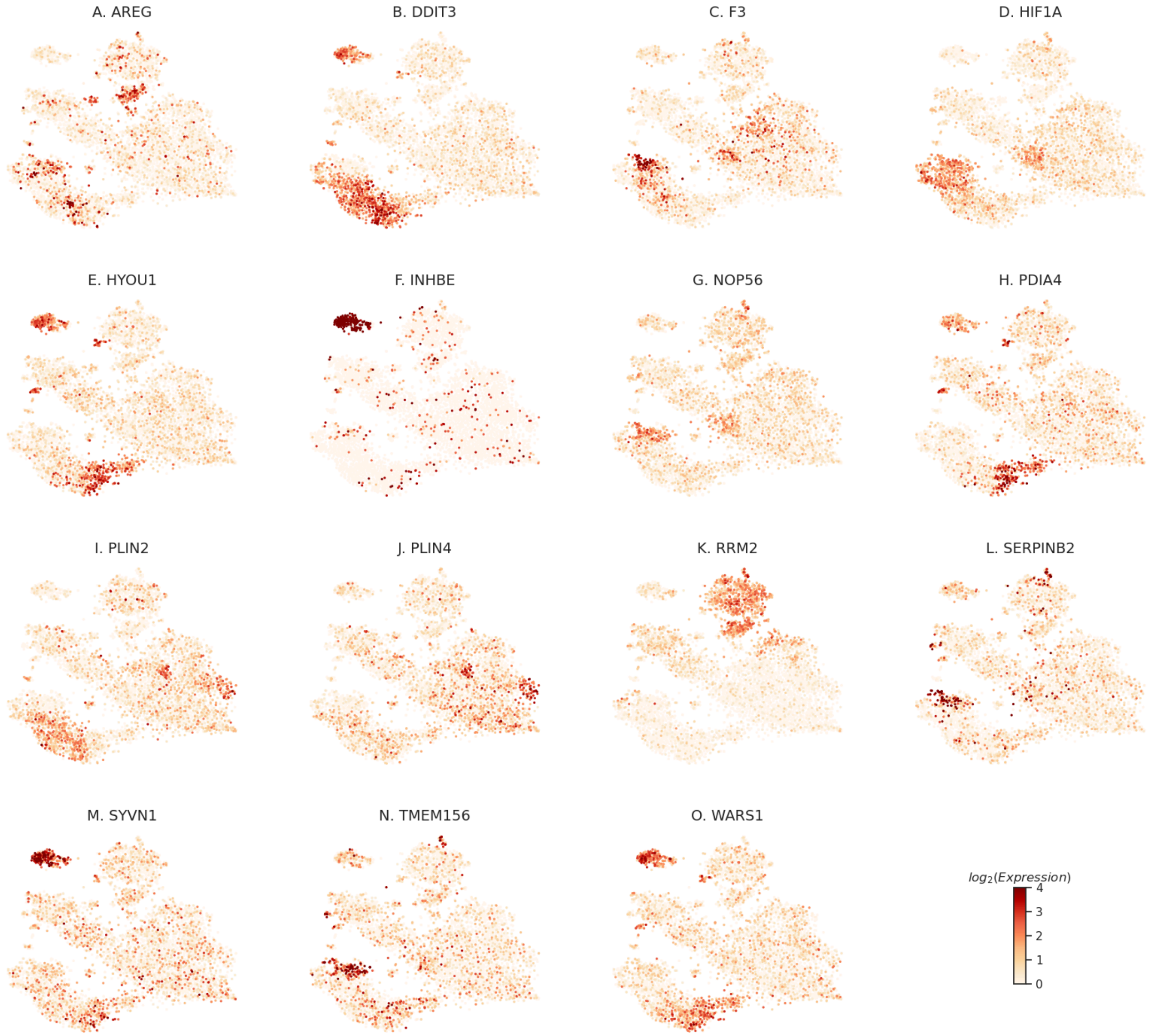


t-SNE plots showing expression patterns of selected genes across single cells. Each panel (A–O) represents one gene, with cells colored by $\log_2$-transformed expression levels (scale: 0–4). A shared colorbar at the bottom right indicates expression intensity. These plots highlight the variability and distribution of gene expression across the cell population.

#### Figure S6: Expression of key cell-type genes across single-cell populations

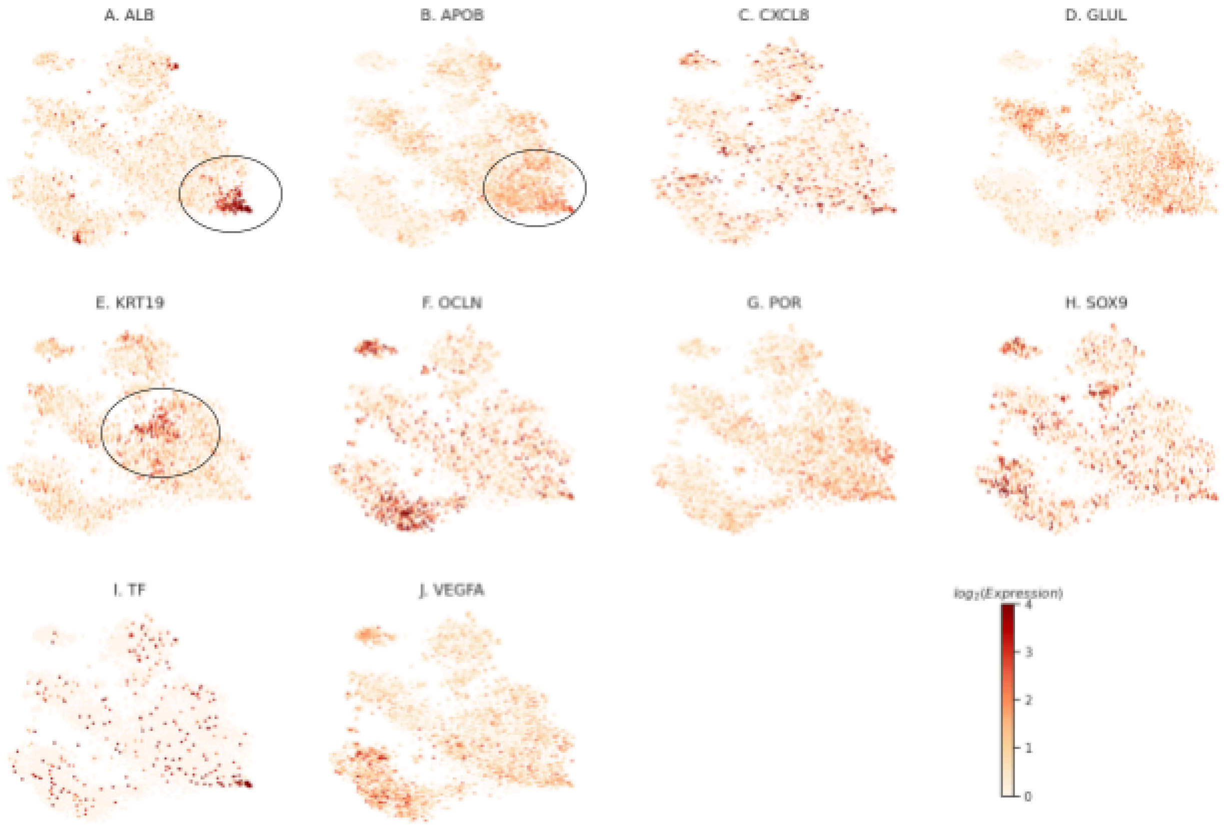


t-SNE plots showing expression patterns of selected cell-type-specific markers across single cells. Each panel represents one gene, with cells colored by $log_2$-transformed expression levels (scale: 0–4). A shared colorbar at the bottom right indicates expression intensity. These plots highlight the variability and distribution of gene expression across the cell population. Figures S6A and S6B show putative subpopulations of hepatocytes, while Figure S6E shows putative cholangiocytes.
